# Timing the initiation of sex: delay mechanisms alter fitness outcomes in a rotifer population model

**DOI:** 10.1101/2025.04.25.650506

**Authors:** Bethany L. F. Stevens, Silke F. van Daalen, Tirzah J. Blomquist, Kristin E. Gribble, Michael G. Neubert

**Affiliations:** Department of Ecology, Evolution, and Marine Biology. University of California Santa Barbara, Santa Barbara, CA, 93106, USA; Wageningen Marine Research, 1976 CP, IJmuiden, NL; Biology Department, Woods Hole Oceanographic Institution, Woods Hole, MA, 02543, USA; Josephine Bay Paul Center for Comparative Molecular Biology and Evolution, Marine Biological Laboratory, Woods Hole, MA, 02543, USA

**Keywords:** Life History Strategy, Environmental Stochasticity, Cyclical Parthenogenesis, Rotifers, Dormancy, Delay Differential Equations

## Abstract

Species that inhabit variable environments have complex mechanisms to precisely time their life-history transitions as conditions change. One such mechanism in rotifers is a block on sexual reproduction that extends across multiple asexual generations after emergence from diapause. It has been hypothesized that this delay is advantageous in competitive and stochastic environments. Here, we develop a model of cyclically parthenogenic rotifer populations with a novel formulation of a “mictic block” that prevents sexual reproduction by females that are not sufficiently distant, generationally, from a stem ancestor that was produced sexually. We find that mictic blocks are indeed adaptive but that the most successful phenotypes have shorter blocks than previously reported and that the success of different delay phenotypes is highly dependent on the duration of the growing season. For a fixed environmental regime, a stable polymorphism is possible, wherein a phenotype with a longer block performs better in years with an average-length growing season, and a phenotype with a shorter block and lower mixis ratio performs better in years with an “extreme” growing season, whether short or long. Our model provides an eco-evolutionary framework for the study of *Brachionus* rotifers, a model system for non-genetic maternal effects and the evolution of sex.

## 1. Introduction

For species that live in variable environments, the timing of life history transitions is often subject to strong selection pressure. Mistiming activities such as metamorphosis, reproduction, or emergence from dormancy relative to changes in the environment can significantly reduce fitness. As a result, species living in variable environments have evolved precise, sometimes elaborate, mechanisms to control the timing of life-history transitions. Many species rely on a combination of external cues, such as photoperiod or food availability (Zhang and Baer, 2000; Koch et al., 2009), and internal timing mechanisms, or biological clocks (Simon et al., 2001). Although external cues may provide information about imminent changes in conditions, internal clocks can protect populations from unreliable cues or abrupt fluctuations.

*Brachionus* rotifers that live in ephemeral ponds are examples of organisms with carefully timed life-history transitions. At the beginning of the growing season, resting eggs in the sediment hatch into asexual (amictic) females that live in the water column (Fig. 1). After a period of relatively rapid population growth fueled by asexual reproduction, amictic females begin producing mictic female offspring capable of sexual reproduction along with amictic female offspring. In the absence of fertilization by males, mictic females produce eggs that develop into males. Those males fertilize mictic females, who then produce resting eggs—stress-resistant, dormant embryos that can survive when the pond dries or freezes between growing seasons (García-Roger et al., 2019).

**Figure 1:**
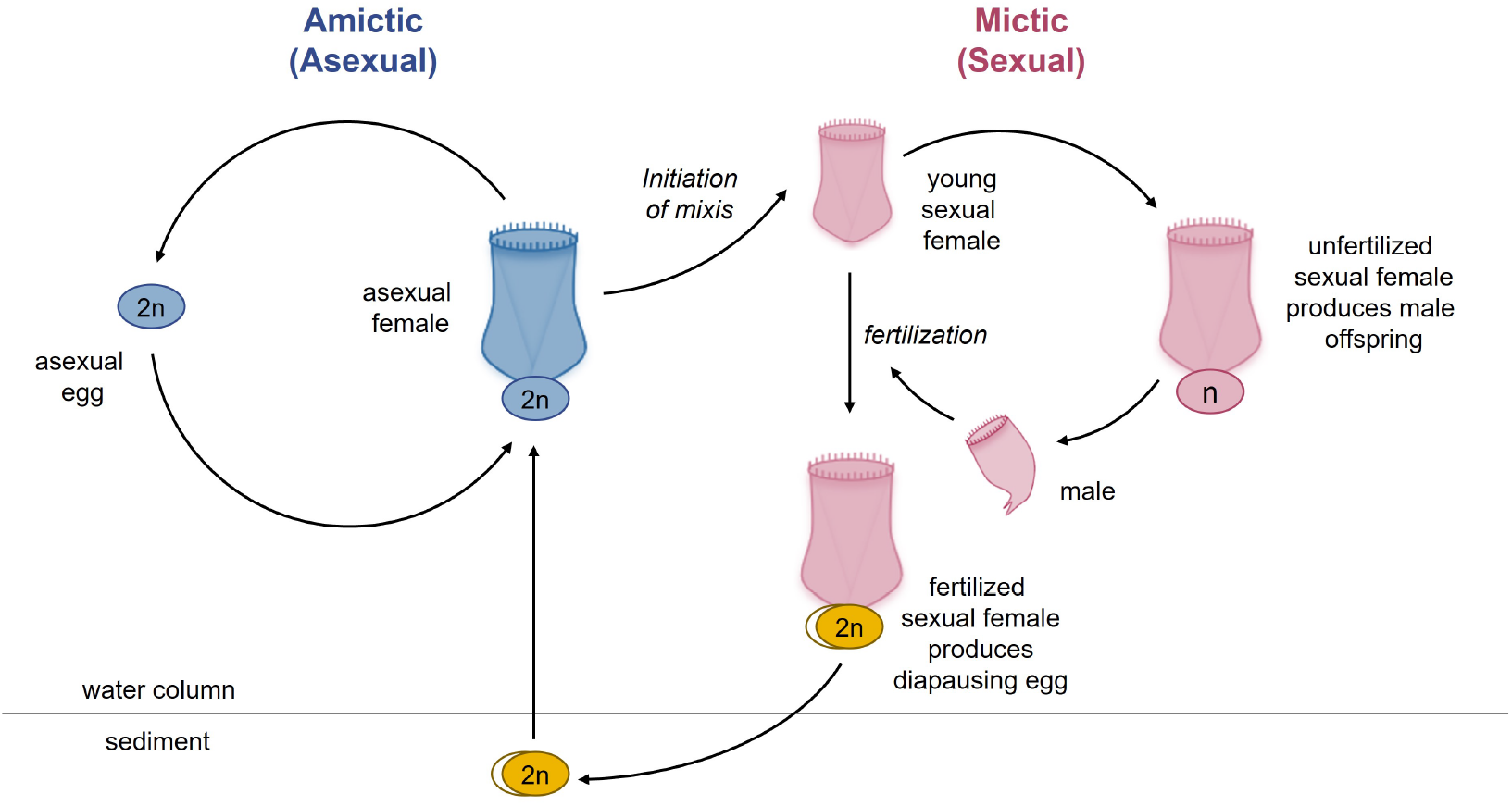
Life cycle of the rotifer *B. manjavacas*. Left (blue), the asexual (amictic) cycle, in which a female produces clonal diploid eggs by mitosis. Right (pink), the sexual (mictic) cycle, in which crowding conditions prompt a portion of females in the population to become mictic, producing haploid gametes via meiosis. If haploid gametes are not fertilized, they hatch into diminutive haploid males. Fertilized gametes develop into dormant resting eggs, able to desiccate and overwinter in the sediments. Upon hatching, resting egg restore the asexual cycle. Induction of the sexual phase is blocked for asexual females who are not sufficiently removed from their last sexual ancestor. Figure and caption adapted from Gribble (2021).

This reproductive strategy, in which several rounds of clonal reproduction are followed by a sexual event, is called *cyclical parthenogenesis*. In animal species, cyclical parthenogenesis commonly occurs in trematode parasites, cladocerans (e.g., *Daphnia* spp.), aphids, cecidomyiids (gall midges) and cynipids (gall wasps), in addition to rotifers (Neiman et al., 2014). The strategy strikes a delicate balance between the contrasting benefits of sexual and asexual reproduction (Bogdan and Gilbert, 1982; Gilbert, 2020). Critical to the success of cyclical parthenogenesis is the *mictic delay* : the time between the start of each growing season and the initiation of mixis, which determines how large the population becomes before sexual reproduction and the production of resting eggs begin.

Empirical work has identified both external and internal timing mechanisms that influence the length of the mictic delay in rotifer populations. Within the genus *Brachionus*, mixis is initiated in response to a density-dependent chemical signal (Carmona et al., 1993; Stelzer and Snell, 2003). In this case, the length of the mictic delay will depend on the time it takes for the rotifer population to exceed a given density, the *mixis threshold density* (Gilbert, 2017). Additionally, Gilbert (2002, 2003) showed that some strains exhibit a secondary, internal delay mechanism. In these populations, individuals do not become receptive to the density-dependent signal until many (8-12) generations after hatching from a resting egg. We refer to this second delay mechanism as a *mictic block*. Internally-timed mictic blocks are hypothesized to be particularly evolutionarily advantageous for late-hatching clones or when a strain is in competition with others that might interfere with its density-dependent chemical signal (Gilbert, 2017, 2020).

Serra et al. (2005) tested this hypothesis with a model of a cyclically parthenogenic rotifer population in a stochastic environment. In their model, females hatch from resting eggs and reproduce asexually until both a mixis threshold density has been met and a minimum number of days have passed since the resting eggs hatched. Serra et al. (2005) used this model to demonstrate that a rotifer phenotype with a mictic block could successfully invade a resident phenotype without such a block, by growing asexually to larger population sizes before transitioning to mixis.

Notably, there is a distinction between a block that lasts a fixed number of days (*temporal block*), as Serra et al. (2005) explored, and a block that lasts a fixed number of generations (*generational block*), as suggested by more recent observations (Colinas, unpublished; section S1; Fig. S2). In the first case, a single individual could become sensitive to the mixis signal over its lifespan, which is inconsistent with current understanding of the rotifer life cycle. Because the timing of the mictic delay is critical to fitness in this system, we must carefully consider how the mictic block is implemented, both biologically and in model formulations.

Here, we investigate how this difference in block implementation alters the fitness and eco-evolutionary dynamics of a cyclically parthenogenic species. We demonstrate that a phenotype with a temporal block produces resting eggs at a higher rate than a phenotype with a comparable generational block (S1). Then, inspired by Serra et al. (2005), we construct a model of a rotifer community with the capacity to delay mixis, in our case for a fixed number of generations. We explore how this new model alters the conclusions of the previous work, and identify fitness-maximizing and evolutionarily stable strategies for timing the onset of mixis.

## 2. Methods

### 2.1. A rotifer model with a generational mictic block

Following Serra and King (1999) and Serra et al. (2005), we model the dynamics of a population of rotifers over the course of sequential growing seasons of variable length (Fig. 2). We take the growing season to be the period of time in which a vernal pool provides viable habitat in the water column, after which the pool dries or freezes; only resting eggs survive to the following growing season. We model the changes in the population densities of females at each life stage over the course of each growing season as follows.

**Figure 2:**
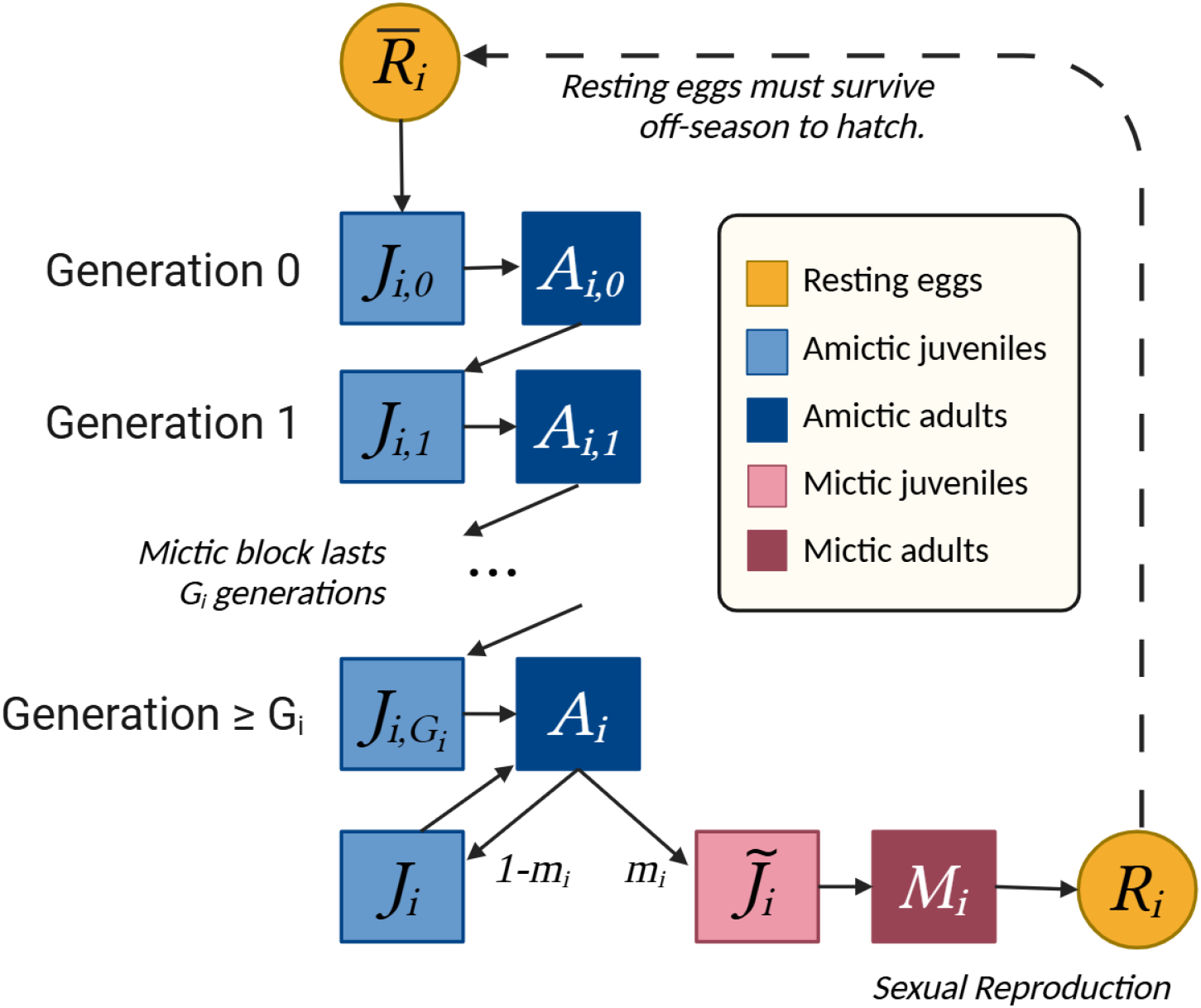
Schematic of cyclically parthenogenic life cycle model with generational block. Resting eggs can only produce stem juveniles (generation 0). Juveniles become adults of the same generation. Amictic adults can subsequently reproduce asexually to create juveniles in the next generation. *G*_*i*_ is the first generation which might produce mictic offspring if other conditions for mixis are met (equation 5). In this diagram, *G*_*i*_ *>* 1, but it could be any nonnegative integer (Fig. S3). Mictic adults only produce males (not included in the model) or resting eggs that will not hatch until the following season.

We begin with an initial density of resting eggs in the sediment that hatch daily in clutches of equal size at times *t* = 0, 1, …, *H* (Serra et al., 2005). Let *ϕ*_*i*_ be the density of eggs of each phenotype *i* divided by *H* + 1, such that every day in this hatching period, *ϕ*_*i*_ stem juvenile females of phenotype *i* are introduced into the water column. Assume that individuals die at rate *q* and that juveniles mature into adults after *τ* days. The population density of stem juveniles of phenotype *i* at time *t*, or *J*_*i*,0_(*t*), therefore changes as

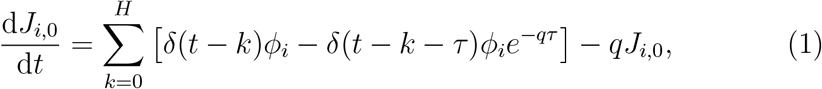

where *t* is time measured in days since the start of the growing season and *δ*(*t*) is the Dirac delta function. The first term within the brackets describes the introduction of clutches that emerge from diapause, while the second term accounts for the maturation of individuals that survive to adulthood *τ* days after they emerge. Maturation leads to the production of mature, amictic stem females of phenotype *i*, which also die at rate *q*:

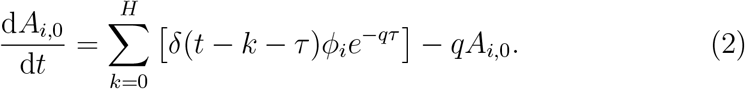

Juvenile females of subsequent generations (*j >* 0) are produced by amictic adults. Asexual reproduction in rotifers has been observed to depend on both crowding and resource availability, with individuals ceasing reproduction in order to survive under starvation conditions (Gatto et al., 1992; Kirk, 1997; Snell et al., 2001). We therefore model birth rate as dependent on the total rotifer density while mortality rate remains density-independent. Let *A*_*i,j*_(*t*) be the population density at time *t* of mature amictic females, with phenotype *i, j* generations removed from a stem female (for *j >* 0). These individuals will produce juveniles of generation *j* + 1 at a per capita birth rate, *b*, which is a function of total population density in the water column, *N* (*t*):

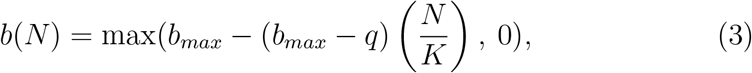

where

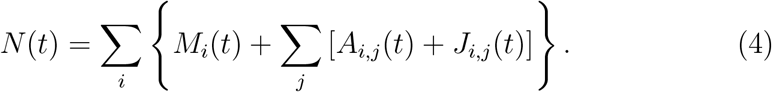

Here, *b*_*max*_ is the intrinsic birth rate, *K* is the community-wide carrying capacity, and *M*_*i*_(*t*) is the population density at time *t* of mature mictic females of phenotype *i*. In this formulation, the birth rate equals the death rate when *N* (*t*) = *K*, although note that if resting eggs continue to hatch, net population growth may not be zero.

In the model, all offspring inherit the phenotype of their mother. The phenotype of each individual comprises three traits: its mixis ratio, *m*_*i*_; mixis threshold density, *T*_*i*_; and the number of generations before mixis can occur, *G*_*i*_. The mixis ratio is the proportion of female juveniles produced by amictic adults that will mature into mictic adults *when mixis is initiated*. The realized mixis ratio, 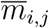, is a function of the individual’s generation (*j*) the population density at the time of reproduction (not at the time of maturation), and the individual’s phenotype (*i*):

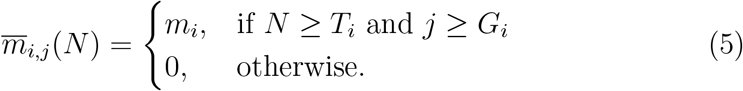

While 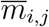 and *m*_*i*_ are strictly proportions (0 *< m*_*i*_ *<* 1), we refer to these terms as mixis ratios, in keeping with the rotifer literature (e.g., Seudre et al., 2020). We explore evolutionary changes in all three dimensions of rotifer phenotype in our model through evolutionary invasion analysis (Section 3.2).

Let the total rate of asexual reproduction by phenotype *i* amictic adults of generation *j* at time *t* be expressed as Ψ_*i,j*_(*t*) = *b*(*N* (*t*))*A*_*i,j*_(*t*). The dynamics of juvenile, amictic adult, and mictic adult females of all generations greater than 0 are then:

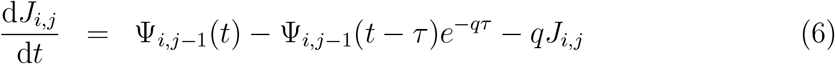

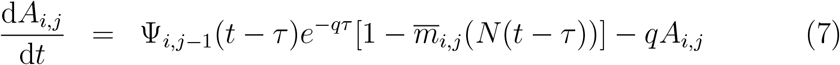

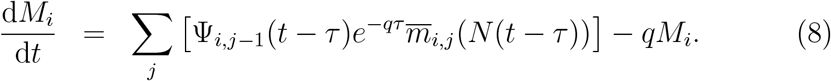

The three terms in equation (6) account for births, maturation of juveniles that survive their juvenile period, and mortality. The first terms in equations (7) and (8) represent the maturation of juveniles into amictic and mictic adults, respectively. We do not distinguish between generations of mictic adults, so the production of mictic adults by all generations is summed.

Finally, we simulate the number of resting eggs produced by mictic females over the course of each growing season. Let *R*_*i*_(*t*) be the population density of the resting eggs produced during the current growing season by phenotype *i* at time *t*. Resting eggs are produced at a reduced rate relative to the rate that amictic females produce juveniles. The rate is reduced by a factor, 0 *< c <* 1, reflecting the costs of sexual reproduction, including the production of males and increased resource allocation to resting eggs. Resting egg production is therefore described by

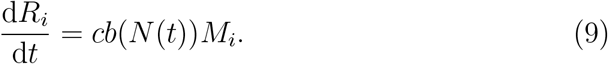

As in Serra et al. (2005), we take the length of each season, *S*, to be a uniformly distributed random variable between *S*_*α*_ and *S*_*ω*_. Thus, *R*_*i*_(*S*) is the density of eggs of phenotype *i* that had been produced by the end of a season. There is a maximum density, *B*_*max*_, of resting eggs across all phenotypes that can survive to hatch in the next growing season. Let 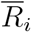 denote the density of eggs of phenotype *i* that survive to the next season. The between-season dynamics are then defined by

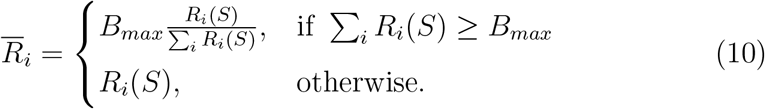

All state variables except 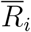 reset to 0 between seasons (since no individuals in the water column survive past the growing season’s end and no new eggs are laid until the next season begins). The first hatching in the new season will produce 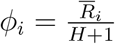 individuals at time *t* = 0.

In our simulations, we calculate the realized fitness of phenotype *i* within each season as the ratio of eggs produced to the number of eggs that initialized the population:

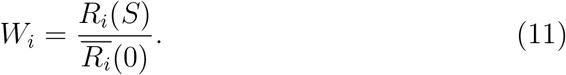

Because the season length *S* varies randomly, so do the the ratios *W*_*i*_ across seasons. We also consider the expected number of resting eggs laid per season, which is the average value of *R*_*i*_(*S*) across all the seasons within a simulation.

A complete list of parameters and default values is included in Table 1. Serra et al. (2005) provide empirical justification for the values of each biological parameter except *B*_*max*_ and *H*. We therefore investigate sensitivity of fitness to these two parameters and to season length.

**Table 1:**
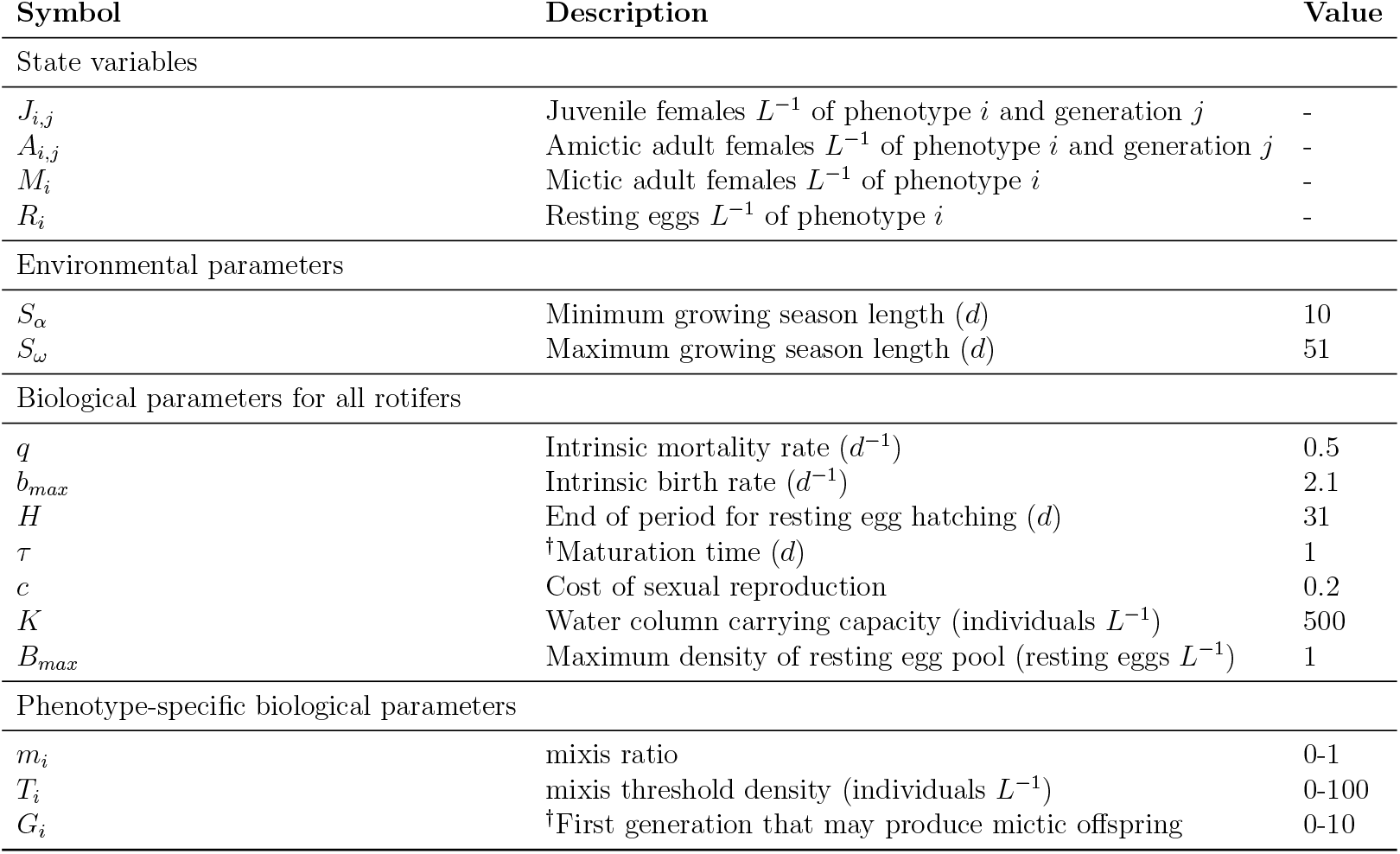
Model state variables and parameters. Daggers in descriptions indicate variables that were added to our model relative to Serra et al. (2005). All parameter values were chosen to match those in previous work, except *b*_*max*_ which was increased to exactly adjust for the lost reproduction from individuals that do not survive their maturation step in our model.

| Symbol | Description | Value |
| --- | --- | --- |
| State variables |  |  |
| $J_{i,j}$ | Juvenile females $L^{-1}$ of phenotype $i$ and generation $j$ | - |
| $A_{i,j}$ | Amictic adult females $L^{-1}$ of phenotype $i$ and generation $j$ | - |
| $M_i$ | Mictic adult females $L^{-1}$ of phenotype $i$ | - |
| $R_i$ | Resting eggs $L^{-1}$ of phenotype $i$ | - |
| Environmental parameters |  |  |
| $S_\alpha$ | Minimum growing season length ($d$) | 10 |
| $S_\omega$ | Maximum growing season length ($d$) | 51 |
| Biological parameters for all rotifers |  |  |
| $q$ | Intrinsic mortality rate ($d^{-1}$) | 0.5 |
| $b_{max}$ | Intrinsic birth rate ($d^{-1}$) | 2.1 |
| $H$ | End of period for resting egg hatching ($d$) | 31 |
| $\tau$ | <sup>†</sup>Maturation time ($d$) | 1 |
| $c$ | Cost of sexual reproduction | 0.2 |
| $K$ | Water column carrying capacity (individuals $L^{-1}$) | 500 |
| $B_{max}$ | Maximum density of resting egg pool (resting eggs $L^{-1}$) | 1 |
| Phenotype-specific biological parameters |  |  |
| $m_i$ | mixis ratio | 0-1 |
| $T_i$ | mixis threshold density (individuals $L^{-1}$) | 0-100 |
| $G_i$ | <sup>†</sup>First generation that may produce mictic offspring | 0-10 |

## 3. Analysis

### 3.1. Monomorphic Populations

We begin our analysis by exploring the dynamics of populations with a single shared phenotype. For these populations, our model produces qualitatively similar dynamics to those of Serra et al. (2005) (Fig. 3 A). Population growth starts slowly as resting eggs gradually hatch to produce stem juveniles and subsequently stem adults. The most rapid growth is achieved while these adults reproduce entirely asexually. Once their phenotype’s mixis threshold density, *T*_*i*_, is reached, all amictic adults that are of generation *G*_*i*_ or greater will begin producing some mictic adults which create resting eggs. The net growth of amictic individuals in the water column declines and the density of resting eggs increases.

**Figure 3:**
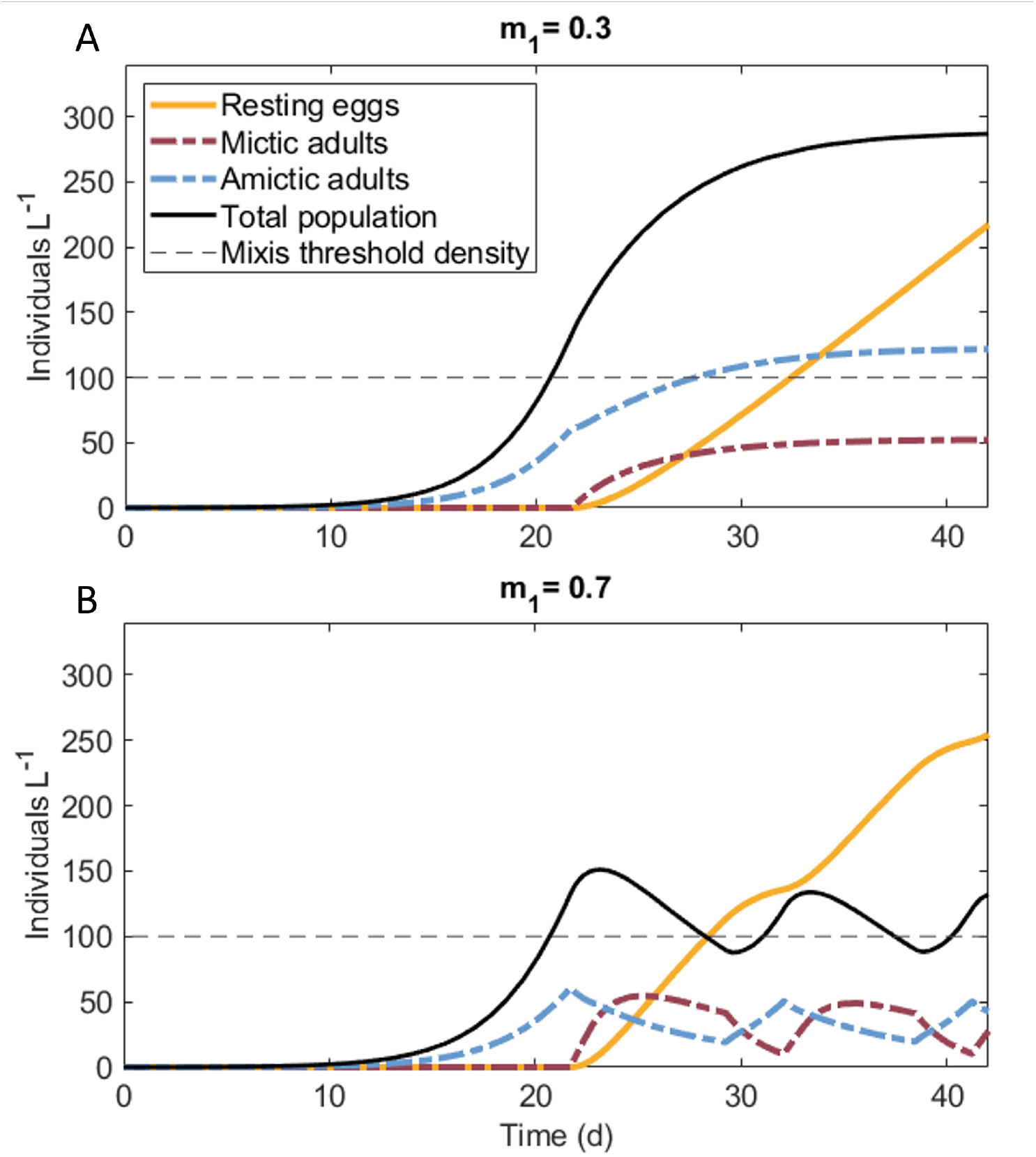
Dynamics within a single season for populations with no generational mictic block (*G*_*i*_ = 0). Parameter values for these simulations are those indicated in Table 1, with initial resting egg density equal to *B*_*max*_, mixis ratio (*m*_1_) phenotypes indicated in the title to each panel, and mixis threshold density indicated by the horizontal dashed line (*T*_1_ = 100). When the mixis ratio is high enough to lead to population decline (B), the total population can oscillate around the mixis threshold. See Fig. S4 for a close-up of hatching and maturation dynamics.

In general, our simulated populations produce resting eggs more slowly than those presented in Serra et al. (2005). As demonstrated in section S1, the generational mictic block, compared to a temporal mictic block, results in a lower rate of resting egg production at the onset of mixis. This exacerbates the risks and reduces the benefits of waiting to initiate mixis. We therefore expect the optimal threshold and block traits to be lower in our analysis compared to those reported in the previous work.

To test this hypothesis, we analyzed our model for the same environmental regime as explored in Serra et al. (2005). That is, we ran simulations for 40 consecutive seasons wherein each season length was randomly chosen from a uniform distribution between 10 and 51 days and assessed differences in average fitness between phenotypes (Fig. 4). Patterns in fitness across phenotypes appear to be insensitive to the values of *H* and *B*_*max*_ (Fig. S7, S8). In the absence of a generational block, we find that the fitness-maximizing phenotype is a mixis ratio of 25% and a threshold value of 0.4 rotifers L^−1^ (for 0 *< m*_*i*_ *<* 1, *T*_*i*_ *<* 100 *L*^−1^; Fig. S5). This contrasts with the values of 45% and 82 rotifers L^−1^ reported in Serra et al. (2005). Looking at individual seasons (Fig. 5), we see that long seasons produce fitness patterns similar to those presented in Serra et al. (2005). When seasons are short, however, phenotypes with high threshold densities fail to produce any resting eggs, which leads to zero fitness. Because of the lower rates of resting egg production in our model, selection pressure favors much lower threshold densities than previously reported for the same environmental regime.

**Figure 4:**
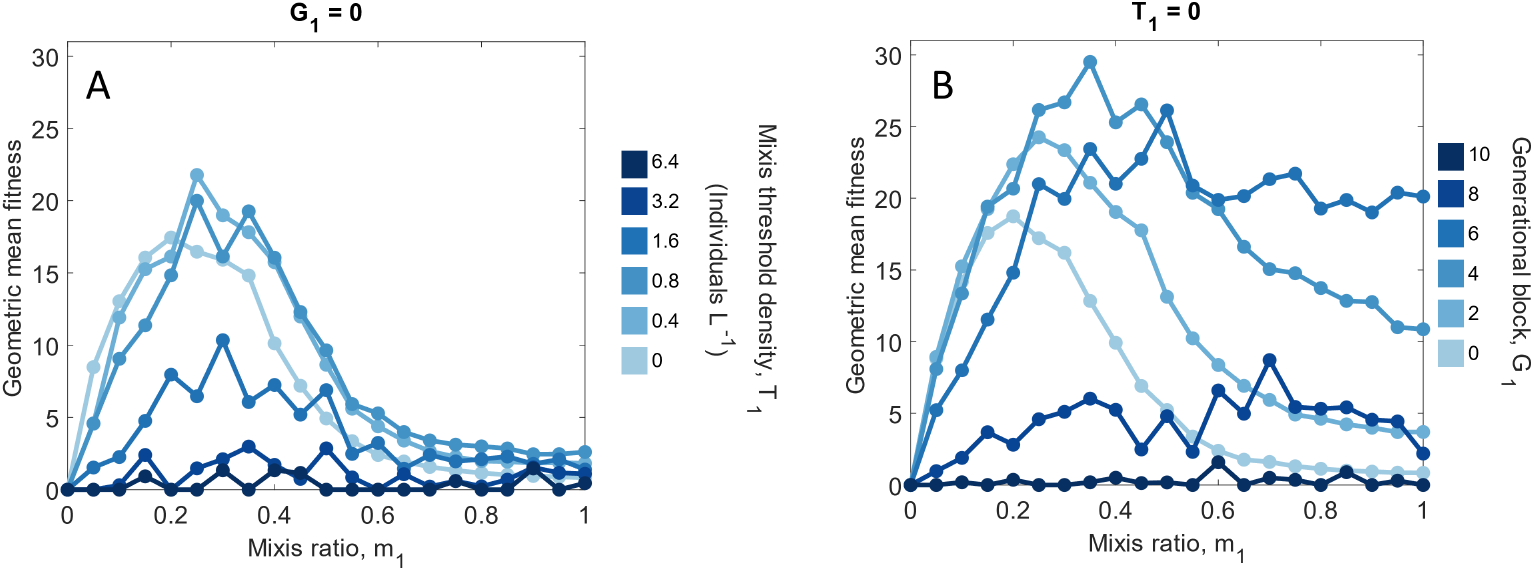
Fitness of monomorphic populations of phenotypes without a generational mictic block (A) and without a mixis threshold density (B). Values plotted are the averages across 40 replicates. Generational mictic blocks of up to 6 generations generally increase fitness. In the absence of a mictic block, phenotypes with low mixis ratios and low mixis threshold densities have highest fitness. All parameters not indicated within the figure have values shown in Table 1. A close-up of panel (A) at a higher resolution is provided as Fig. S5, and a version of this figure with expected egg production per season is included as Fig. S6 .

**Figure 5:**
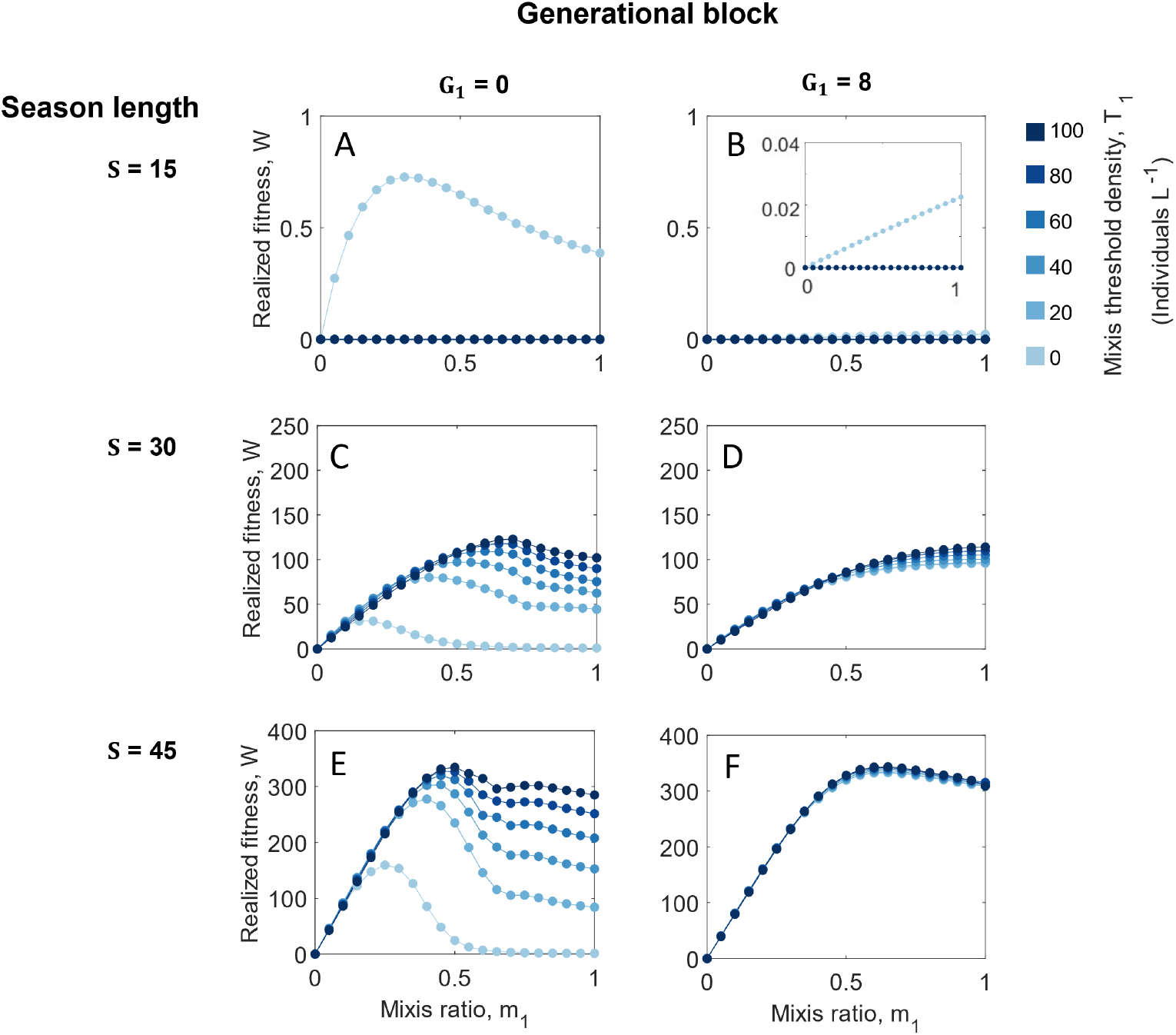
Realized fitness (resting eggs produced per egg hatched) of monomorphic populations within short (A-B), medium (C-D) and long (E-F) seasons of fixed length. Phenotypes with low mixis threshold densities have higher fitness in short seasons, and those with high threshold densities have higher fitness in long seasons. A long generational block (right column) does poorly in short seasons but can increase the fitness of phenotypes with low threshold densities in long seasons. Resting egg production in (B) is low but nonzero (∼ 0.01 *L*^−1^) for phenotypes with *T*_*i*_ = 0. Note that the vertical scale changes between rows. Simulations were run with parameter values indicated in Table 1 and initial resting egg density, *B*_*max*_.

Phenotypes with low threshold densities have the highest fitness when paired with lower mixis ratios (Fig. 4A, Fig. 5C). A phenotype with a low threshold and low mixis ratio will have an early onset of mixis and low rates of sexual reproduction throughout the season. Compared to phenotypes with high mixis threshold densities, one with a low threshold density will have greater fitness in in short seasons and lower fitness in long seasons. Without a generational block, surviving short seasons comes at the cost of reduced fitness in long seasons.

Generational blocks evidently reduce the trade-off between survival in short seasons and high fecundity in long seasons (Fig. 5). Compared to mixis threshold densities, which apply to all cohorts equally, generational blocks are more forgiving in very short seasons. The first cohort of eggs to hatch will lead to individuals of generation *G*_*i*_ after *τ* ∗ *G*_*i*_ days, such that some resting eggs may be produced at that time, even if the cohort is small. At the other end of the spectrum, generational blocks are also successful at producing large numbers of resting eggs in long seasons. Phenotypes with generational blocks and low mixis threshold densities generate roughly the same number of resting eggs as those with higher mixis threshold densities (Fig. 5E-F). We expect this insensitivity to total population density to be particularly advantageous in a competitive context when the abundance of a competing phenotype would interfere with the density-dependent timing cue.

### 3.2. Evolutionary Invasion Approach

We now consider competition between distinct rotifer phenotypes to ask whether a generational mictic block can be evolutionarily advantageous. To answer this question, we employ adaptive dynamics, also called evolutionary invasion analysis (Geritz et al., 1998). This approach explores the scenario in which a small mutation arises among a subset of individuals within a resident population at its asymptotic attractor (e.g., an equilibrium point or limit cycle). The mutant phenotype, or invader, will either increase in frequency in the population or will fail to invade, possibly going extinct. In a deterministic model, these outcomes reflect high and low invasion fitness, respectively, and, in some simple cases, an analytical expression for invasion fitness can be written explicitly. Due to the high-dimensional stage structure, nonlinearities, and stochasticity in our model, we cannot attain an analytical expression for invasion fitness. Instead, we use a numerical estimate of invasion fitness. Note however, that stochasticity can lead to extinction even when a phenotype’s invasion fitness is high. We therefore also conduct numerical invasion experiments in which the complete dynamics of two interacting phenotypes are simulated as in Serra et al. (2005).

We estimate invasion fitness by computing the relative fitness of an invader within a monomorphic resident population. More specifically, we simulate the dynamics of a resident population for a sequence of 1000 seasons, such that the number of eggs produced approaches a stable distribution. We calculate its fitness within each season according to Eq. 11 and consider the geometric mean across all seasons. We then simulate the addition of an invader at low frequencies at the start of each season. The invader frequency is assumed to be so low that it has no impact on the resident dynamics or the total rotifer density. We again compute fitness according to Eq. 11. The relative invasion fitness estimate is taken to be the ratio of the mean fitness of the invader to that of the resident.

To confirm that this estimate of relative invasion fitness corresponds to differences in realized invasion outcomes, we conduct simulated competitive invasion experiments. For each replicate of these experiments, we introduce an invader phenotype into a monomorphic resident population and simulate the competitive dynamics for 40 growing seasons. We initialize the population with *B*_*max*_ resting eggs and an invader frequency of 0.05, and, as in Serra et al. (2005), consider an invasion successful if the invader frequency increases on average over 20 experiments. We conduct invasion experiments across a range of values for both the resident and invader phenotypes, generating pairwise invasibility plots which indicate the long term evolutionary behavior of the model.

One of the pillars of adaptive dynamics is that mutations are assumed to be rare, such that the population reaches its equilibrium between each new mutation. By this logic, we initially assume that invaders differ from the residents in only one of the three dimensions of their phenotypes (*G*_*i*_, *T*_*i*_, and *m*_*i*_) at a time. As in Serra et al. (2005), we first find a stable mixis strategy in the absence of a mictic block (*G*_*i*_ = 0), then allow the length of the mictic block to evolve. Next, we extend our analysis beyond that that of Serra et al. (2005) by considering competition between phenotypes with differences in any of the three distinct mixis traits (*G*_*i*_, *T*_*i*_, and *m*_*i*_). We simulate a series of mutations by conducting invasion experiments in each of the dimensions of the phenotype, where the trait values of the resident are updated before each subsequent invasion. In this way, we arrive at a phenotype that cannot be invaded and replaced by any single trait mutation. Finally, we conduct pairwise invasion experiments between phenotypes that may differ in both *m*_*i*_ and *G*_*i*_ to further explore the stability of the system and the potential for stable polymorphisms.

### 3.3. Results of Evolutionary Invasion Analysis

We find that a mixis ratio of 0.11 is evolutionarily stable among phenotypes without a mictic block or mixis threshold density (0 *< m*_*i*_ *<* 1 at a resolution of 0.01; Fig. 6A). That is, this phenotype can successfully invade and resist invasion by a strain with any other mixis ratio. Note that this value is less than the mixis ratio value that maximizes fitness or resting egg production in the absence of competition (Figs. 4, S6). This result is qualitatively comparable to that reported in Serra et al. (2005), in that the optimal mixis ratio is lower in competition than in isolation.

**Figure 6:**
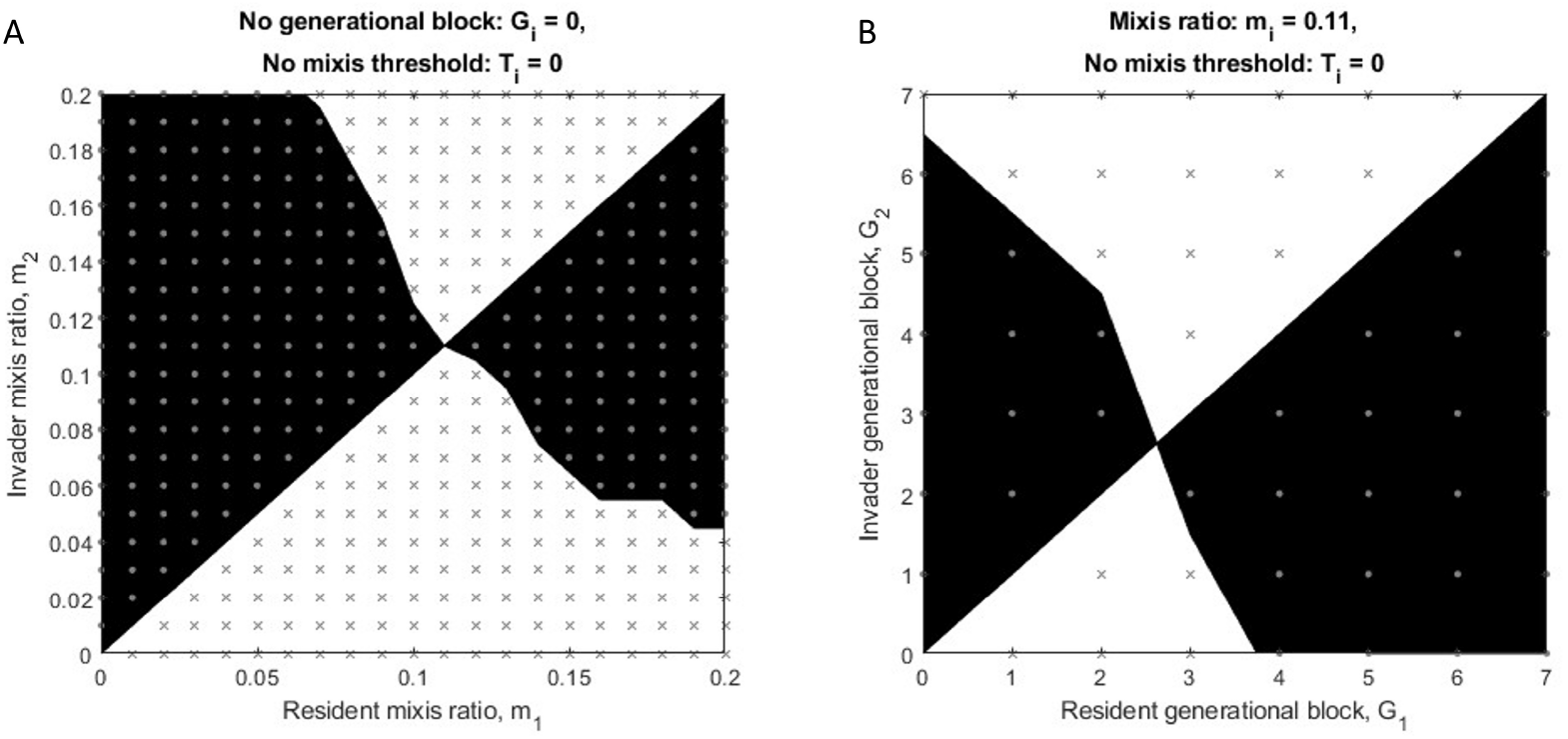
Pairwise invasibility plot for rotifers with A) variable mixis ratio given no generational block or mixis threshold density (*G*_*i*_ = 0, *T*_*i*_ = 0). Invasion was deemed successful if, out of 20 simulations of 40 growing seasons each, the invader frequency increased on average from its initial frequency of 0.05. Gray crosses indicate unsuccessful invasions; circles indicate successful invasions. Black and white regions are interpolated regions of success and failure, respectively. Without a mixis block, a mixis ratio of 0.11 is the best invader and most resistant to invasion. B) Pairwise invasibility plot for rotifers with variable generational blocks given a mixis ratio of 0.11 and no mixis threshold density. Phenotypes with *G*_*i*_ = 3 are most resistant to invasion and can successfully invade all others. All parameters not indicated within the figure have values shown in Table 1.

In general, we find that residents are susceptible to invasion by phenotypes with lower mixis threshold densities. Among phenotypes without a mictic block and with a mixis ratio of 0.11, phenotypes without a mixis threshold density can successfully invade strains with any other value of *T*_*i*_ (Fig. S9A). Competitive outcomes of phenotypes that differ in *T*_*i*_, however, are sensitive to stochasticity in season length, creating noise in our pairwise invasibility plots. Nonetheless, phenotypes without mixis thresholds were successful invaders across all the phenotypes we tested.

We next ask whether a rotifer phenotype that delays mixis through a generational block can invade the strategy that would be evolutionarily stable in the absence of a block. That is, we allow *G*_*i*_ to mutate in a resident population with no block, a mixis threshold density of 0, and a mixis ratio of 0.11. We find that phenotypes with blocks that last 2 or 3 generations have high invasion fitness and can regularly invade and replace phenotypes without a block (Fig. 7). Phenotypes with blocks of 4 generations also have relatively high fitness, but their actual invasion success is much more sensitive to the stochasticity in season length. Blocks longer than 5 generations are mostly unsuccessful (Fig. 7).

**Figure 7:**
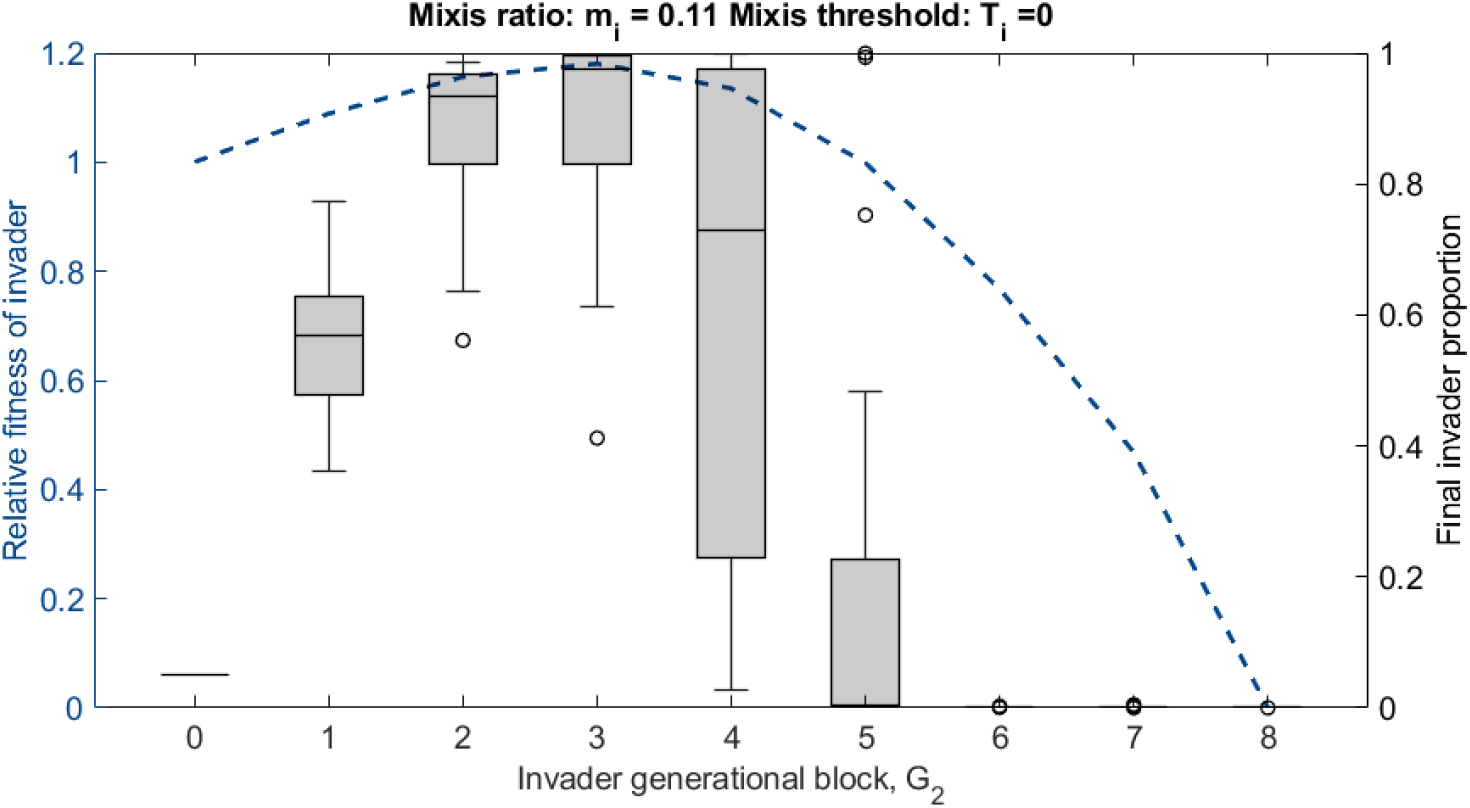
Relative fitness and invasion success of phenotypes with generational mictic blocks. Invaders were introduced into a resident population with the most competitive mixis ratio and threshold phenotype in the absence of a mictic block (*m*_*i*_ = 0.11, *T*_*i*_ = 0). Strains with this mixis strategy and generational blocks of 3 generations have the highest relative fitness and can regularly invade and replace a resident without a block. Each box indicates the median and interquartile range (IQR) of 20 simulations of 40 seasons each per phenotype. Circles indicate values more than 1.5 ∗ IQR away from the nearest box edge. All other parameters for these simulations are those indicated in Table 1.

Our results contrast quantitatively with the findings of Serra et al. (2005), which identified the optimal invader block timing as 8-12 days, substantially longer than the three days it would take to produce adults of generation 2. The difference is in line with our expectations described in section S1. The temporal block of Serra et al. (2005) results in a higher per capita rate of egg production after the block is lifted, artificially favoring a longer delay and a higher mixis ratio. In contrast, our generational block implementation favors a shorter mictic delay to make up for the slower production of resting eggs. Even with the more realistic generational block model formulation, we find that a block phenotype is adaptive, validating the conclusions of Serra et al. (2005). We note that the invasion outcomes for any specific block length depends critically on the environmental regime and the relative chances of experiencing a short or a long season.

We now extend our invasion experiments to explore mutations in any of the three mixis traits. Starting with the phenotype that had the highest fitness in the absence of a generational block (*G*_*i*_ = 0, *T*_*i*_ = 0.4, *m*_*i*_ = 0.25), we conduct a series of pairwise invasibility experiments, each of which simulates evolution along a single phenotype dimension at a time. We update the default phenotype of the resident to reflect the strategy that is evolutionarily stable along that single dimension before conducting a new pairwise invasibility experiment along a different trait axis (Fig. S10). After eight iterations, we find that the simulated population has reached a phenotype (*m*_*i*_ = 0.17, *T*_*i*_ = 0, and *G*_*i*_ = 3) that cannot be replaced by phenotypes with mutations in any single trait.

Lastly, we conduct a collection of invasion experiments in which residents and invaders vary in both their values of *G*_*i*_ and *m*_*i*_ (Fig. S12). We find that for all phenotypes within the range of 0 ≤ *G*_*i*_ ≤ 5 and 0.05 ≤ *m*_*i*_ ≤ 0.25, two combinations of traits produce phenotypes that maximize both their invasion success and their resistance to invasion. The first is that which we identified by varying each trait in isolation: *G*_*i*_ = 3, *m*_*i*_ = 0.17. The second has a longer delay and a higher mixis ratio, *G*_*i*_ = 4, *m*_*i*_ = 0.20. When strains with phenotypes in each of these regions compete, each is able to invade the other when introduced at low abundances, indicating the potential for a stable polymorphism. The phenotype with the shorter mictic block and lower mixis ratio does well both in short seasons, due to its early transition to sexual reproduction, and in long seasons, due to its sustained high population densities. In contrast, a phenotype with a longer mictic block and higher mixis ratio can out-compete the first phenotype in seasons of roughly average length (Fig. S13). Competitive outcomes, in this case, depend on the duration of each growing season. Given the fixed environmental parameters on which we have focused in this work, we find these phenotypes can coexist across many seasons.

## 4. Discussion

Cyclic parthenogenesis occurs in at least seven groups of animals, including the best studied cases of rotifers, cladocerans, and aphids (Hebert, 1987). In these systems, there is strong selection pressure on the timing of sex, such that the onset of sex has evolved to be influenced by both internal and external cues (Simon et al., 2002; Zhang and Baer, 2000; Koch et al., 2009). Here, we have developed a model of the generational block observed in rotifers that delays the onset of mixis. This model allows us to investigate the fitness outcomes of rotifer phenotypes with distinct combinations of internal and external timing mechanisms, addressing questions about when different reproductive strategies are most advantageous.

We found that a phenotype with a generational mictic block can successfully invade a resident population that would be evolutionarily stable in the absence of a block (Fig. 6). This result supports the widespread hypothesis that generational blocks are adaptive and confirms the conclusions of Serra et al. (2005), despite the difference in the formulation of the mictic block between our model and theirs (highlighted in Section S1). Generational blocks increase the cumulative number of resting eggs produced in long seasons while still allowing for a minimal number of eggs in short seasons.

We found the optimal length of the mictic block in generations to be shorter than that reported in Serra et al. (2005), but we note that the success of phenotypes with generational blocks depends sensitively on the distribution of season lengths. Long generational blocks offer a larger benefit in longer growing seasons and have a fitness cost when seasons are short. This is consistent with patterns in block phenotypes observed in natural populations, although there is variability among species (Schröder and Gilbert, 2004). Generational blocks are not evident in populations in the Chihuahuan desert, for example, where vernal ponds only last a few days (Schröder et al., 2007). Across eastern Spain, Colinas et al. (2023) found that ponds with longer hydroperiods are home to rotifer strains with longer generational blocks. Those authors and others have attributed the success of block phenotypes to the increased growth of late-hatching clones, though our model indicates that early-hatching clones also benefit from a mictic block in long seasons (Fig. S14). Phenotypes with high mixis ratios and a long mictic delay produce more eggs due to the prolonged period of asexual population growth.

Rotifer populations can exhibit a prolonged mictic delay as a result of either a high mixis threshold density or a generational block. In our model, the fitness of a phenotype with a long generational block is relatively insensitive to its mixis threshold density (Fig. 5), suggesting that either one of the two delay mechanisms is sufficient for high egg production in long seasons. Compared to phenotypes with high mixis threshold densities, those with generational blocks are stronger competitors and produce more resting eggs across a range of season lengths. Given these advantages, we expect generational blocks to be a more prevalent mechanism for timing the initiation of sex, yet empirical work has reported a wide range of mixis threshold densities (2.3 females *L*^−1^ (King and Snell, 1980); 6.6-22.9 individuals *L*^−1^ (Carmona et al., 1995); 147 individuals *L*^−1^ (Snell and Boyer, 1988)). The prevalence of these mixis threshold densities suggests that they have benefits in addition to the associated delays in the initiation of sex.

Possible benefits of high threshold densities include an increased likelihood of sexual reproduction once high population densities have been reached. In this way, a high threshold density would be a strategy to avoid a mate-finding allee effect, which is not typically seen in rotifers but has been shown to be relevant for other cyclically parthenogenic species (Drake, 2004). Alternatively, high densities may anticipate impending resource limitation (Gilbert, 1967). The transition to sexual reproduction would then occur before individuals are likely to experience starvation. While our model does not include any benefits of mixis threshold densities other than the resulting mictic delay, these benefits could be incorporated in models for future work.

Our results also contrast with observations from Seudre et al. (2020), who found that naturally occurring genotypes with the lowest mixis threshold densities also had the highest mixis ratios. This trend is the opposite of what we see in the model (Fig. 5C), reinforcing the hypothesis that high mixis threshold densities are not only a strategy for timing the onset of sex. Rather, they may reflect a difference in the overall investment of a phenotype in sexual versus asexual reproduction.

Because demographic and environmental parameters vary between specific rotifer strains and habitats, we consider the results of our analysis to be qualitative rather than quantitative. Additionally, we acknowledge that there are a number of simplifying assumptions in our model formulation. There may be demographic effects of population density other than negative density-dependent birth rate that we have overlooked. Maturation time, for example, may depend on resource availability, and under some conditions, rotifer survival has been reported to increase under crowding conditions, likely due to a tradeoff with reproduction (Gatto et al., 1992). More pertinent to our research questions, the density dependent mortality of resting eggs between seasons in our model is quite rigid. In real systems, there is unlikely to be a fixed maximum number of eggs that can survive, and resting eggs may be able to survive multiple off-seasons. Survival depends upon the length of the off-season, pond conditions, and predator dynamics, all excluded from our simulations (Hairston Jr. et al., 1995; García-Roger et al., 2006).

That said, our model produces realistic within-season trajectories of both population size and the proportions of mictic females expected in the population. We note that for certain parameter values, our model exhibits oscillatory dynamics that were not evident in previous model formulations (Fig. 3B). When birth rates are low and the mixis ratio is high, the initiation of mixis will lead to population decline. If the population then drops below the mixis threshold density, it will oscillate around *T*_*i*_, toggling between producing mictic offspring and not. Evidence for such an oscillation has been reported in at least one natural population. While monitoring individual strains over time, Carmona et al. (1995) saw that the proportion of mictic females decreased during the growing season after an initial increase, leading to population growth and a second peak in population density in the latter half of the season. Our model indicates that this dynamic could be the result of the time lag between the conditions experienced by a female, which determine the mixis ratio of her offspring, and the contribution of those offspring to population growth once they mature. This is a non-genetic maternal effect, a phenomenon for which rotifers have become an important study system (Seudre et al., 2020; Liguori et al., 2024). Because our model tracks overlapping generations in the rotifer population, we expect it to be a valuable tool for future work on multigenerational maternal effects in this system.

Finally, we have used our model to confirm the conjecture of Serra et al. (2005) that a stable polymorphism is possible within the environmental regime considered in their work. We find that there exist distinct mixis phenotypes that can mutually invade each other. Long-term coexistence, however, would depend upon the distribution of growing season durations from year to year. In real systems, multiple strains of rotifers can exhibit different temporal mixis patterns within the same habitats, including some that are more continuous and others that are more periodic (Carmona et al., 1995). Our simulations suggest that a change in the distribution of season lengths, as would result from changes in rainfall or evaporation, would lead to changes in the relative fitness of these two strategies. We note that while environmental stochasticity (in the form of random growing season lengths) is critical for the formation of a stable polymorphism in the model we analyzed, such life-history polymorphisms can also evolve in constant environments (Landi et al., 2018).

We have argued in this manuscript for the careful consideration of the mechanisms by which life history transitions are timed. For cyclically parthenogenic rotifers, the success of a mictic delay strategy depends not only on the length of the delay, but also on how it is determined in a population. We and Serra et al. (2005) have explored two implementations, but some species may exhibit a mixture of timing mechanisms, including generational blocks as well as effects of birth order or time within the season. We have shown that a generational mictic block can be a robust and effective strategy for timing sexual reproduction in variable environments.

## Data Availability

The code used to carry out this research is publicly available in a GitHub repository. https://github.com/beefowler/Rotifer_GenerationalBlock

## Acknowledgments

We would like to thank Manuel Serra and his collaborators for developing the original questions and modeling approach that have inspired this manuscript and for welcoming our modifications to their work. We would also like to thank Manuel Serra, Stephen Proulx, and our anonymous reviewers for providing helpful comments on the manuscript. BLFS was supported by LS-FMME-00008380 from the Simons Foundation. KEG was supported by R01AG076592 from the National Institute on Aging and by NSF CAREER 1942606.

## Supplementary Information

### S1. Distinguishing between two block implementations

We here present a simplified example to illustrate the difference between the effect of a temporal block on resting egg production and a generational block of the same duration, *τ* .

Consider a population of adults that may either be of the “stem” generation (those that initiate the population at time *t* = 0), *x*_0_(*t*), or of subsequent generations, *x*_1_(*t*). The total population of adults at time *t* is then, *x*_0_(*t*) + *x*_1_(*t*).

All individuals in the population take *τ* days to mature to adulthood, are subject to mortality at the per capita rate *q*, and once mature, produce offspring at the rate 1 per time. If the initial size of the stem adult population is *x*_0_(0) = *X*_0_, then their population size thereafter decays exponentially:

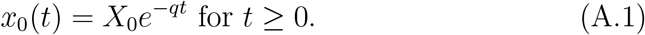

Meanwhile, the population of adults of subsequent (“non-stem”) generations changes according to

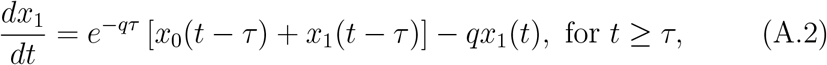

with

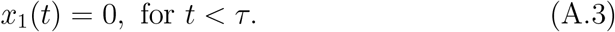

The first term on the right-hand side of equation (A.2) represents the maturation of individuals that survive to adulthood (born to stem or non-stem parents) and the second term describes the mortality of non-stem adults.

Now consider two scenarios in both of which this population generates resting eggs at a per capita rate *r >* 0. In the first scenario, resting eggs are produced only by non-stem adults. This is a generational block of length 1 generation. In this case, the cumulative number of eggs produced by the population up to time *t* will be

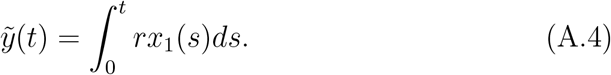

Because no non-stem individuals reach adulthood before time *τ*, we also have

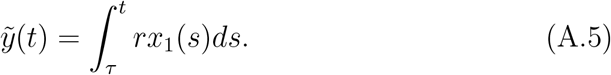

In the second scenario, all adults in the population produce resting eggs after a temporal block equal in length to the maturation time *τ* . In this second case, the number of resting eggs produced will be

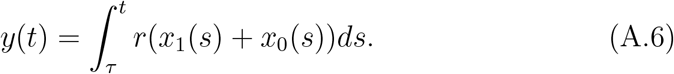

Substituting equation (A.1) for *x*_0_(*s*), equation (A.6) can be rewritten as A.6

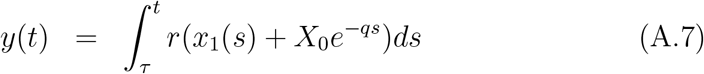

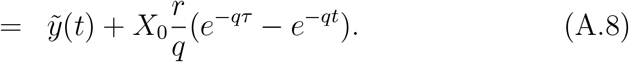

The second term in equation (A.8) is positive for all *t > τ*, so 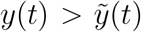. That is, a temporal block produces more resting eggs than a comparable generational block for all *t* (Fig. S1A,B). In both scenarios, no resting eggs are produced until *t* = *τ*, when either the length of the temporal block has passed or the first non-stem adults mature. Yet, the temporal block results in a faster rate of production once the block is lifted. The difference between *y*(*t*) and 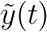 will vary with the initial size of the stem generation, *X*_0_, the egg production rate, *r*, and the duration of the block, *τ* . If resting eggs form the initial population for subsequent growing seasons, egg production by a time-blocked population becomes exponentially larger than egg production by a generation-blocked population over consecutive seasons (Fig. S1C). These differences led us to construct a rotifer population growth model with a generational mictic block, the outputs of which we compare with the outputs of the temporal block model of Serra et al. (2005).

**Figure S1:**
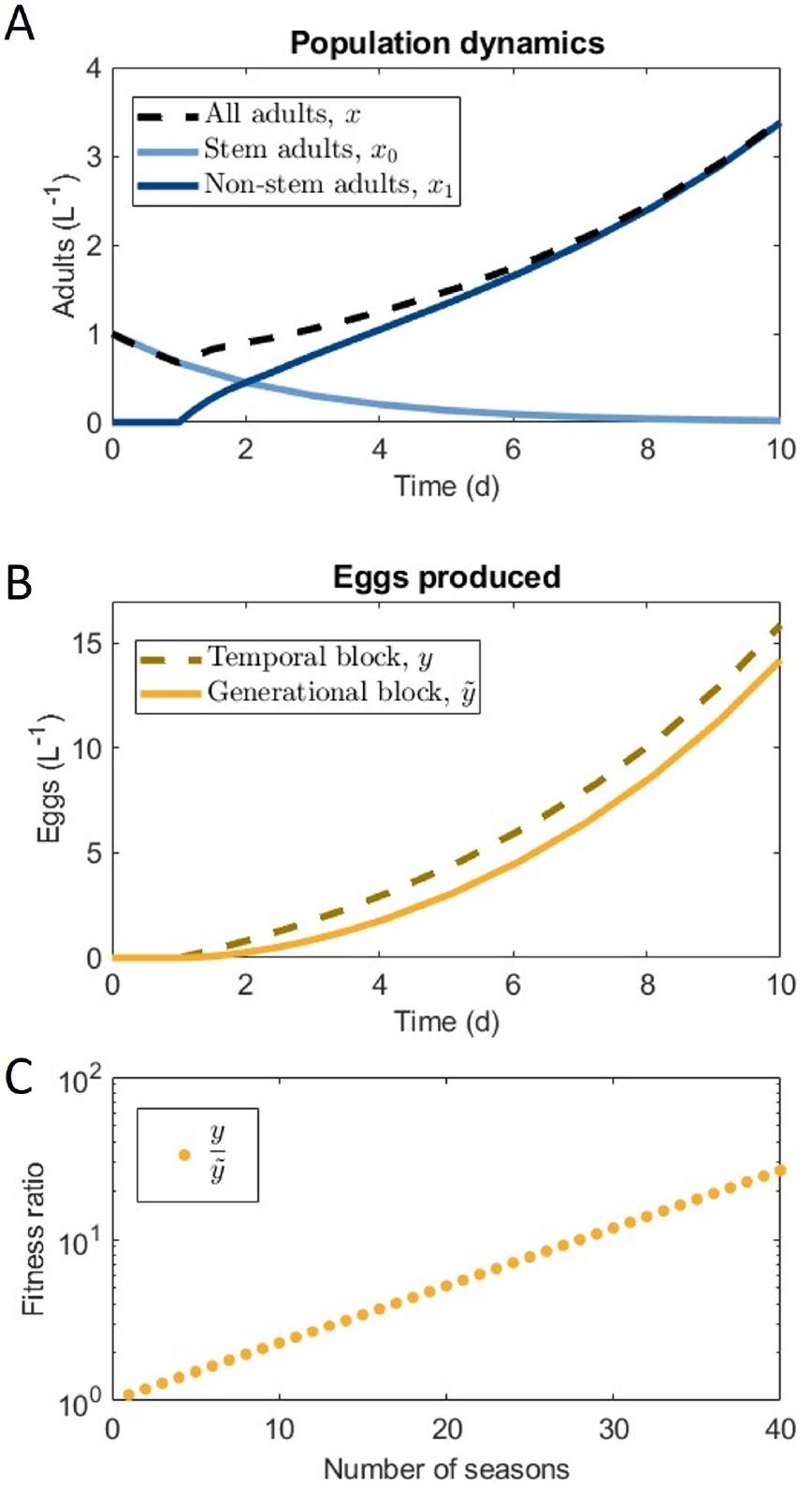
Difference in egg production between phenotypes with a temporal block and a generational block. (A) The number of adults in the population changes according to the model described in equations (A.1-A.3). (B) The number of eggs produced differs between the two different block implementations and depends on the initial size of the stem generation, the egg production rate, and the length of the block; however, the number of eggs produced in the temporal block scenario (dashed) is always greater than that in the generational block scenario (solid). (C) The fitness ratio between the two strategies increases exponentially with the number of growing seasons. Parameter values for these simulations are *X*_0_ = 1, *τ* = 1, and *q* = 0.4. Season length is 10 days.

## Supplementary Figures

**Figure S2:**
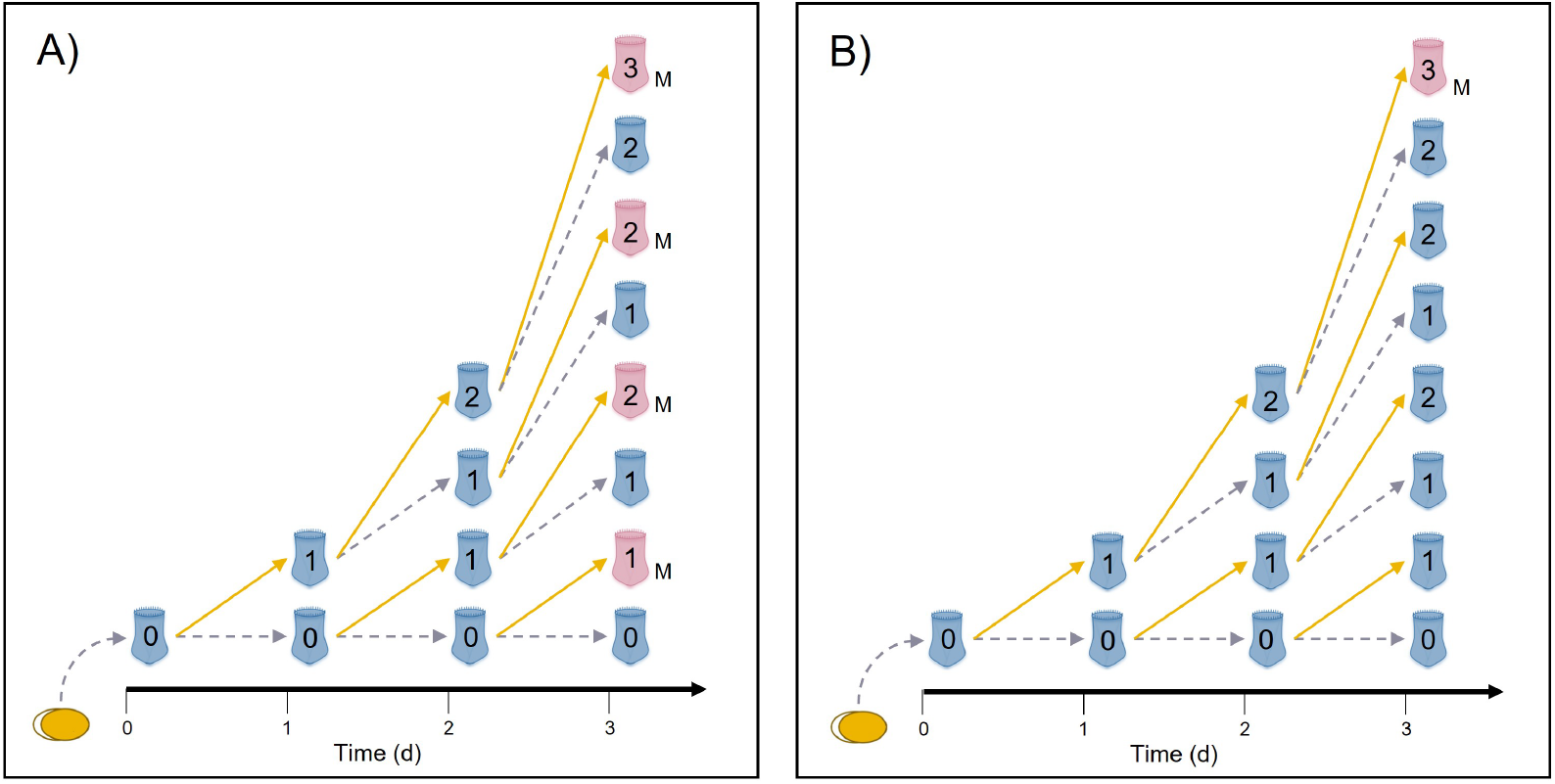
Example population where all rotifers produce one offspring each day demonstrating the difference between (A) a temporal block that lasts two days and (B) a generational block that lasts two generations. An initial resting egg hatches from diapause at *t* = 0. Gray dashed arrows indicate survival and yellow solid arrows indicate reproduction. The number overlaid on each individual indicates how many generations it is removed from the resting egg. For the sake of demonstration, we assume no maturation time, no mixis threshold density, and a mixis ratio of 1. In case (A) the block ends at *t* = 2 and all subsequently produced rotifers are mictic, indicated with an “M”. In (B) only amictic individuals of generation 2 can produce mictic offspring. At day three, there are four mictic individuals in (A) and only one in (B).

**Figure S3:**
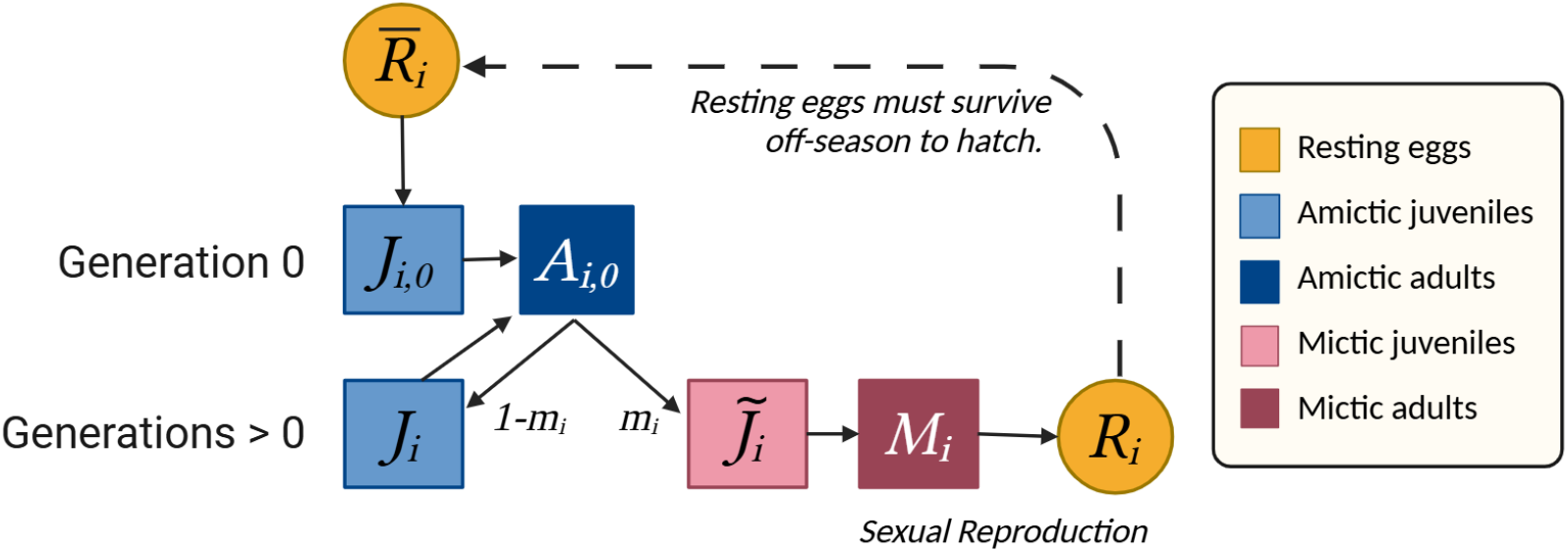
Version of model schematic in Fig., 2, but with *G*_*i*_ = 0. This is the case where the rotifer population has no mictic block. Note that there is still a mictic delay between resting egg hatching and new resting egg production that consists of the maturation time of stem and then mictic juveniles.

**Figure S4:**
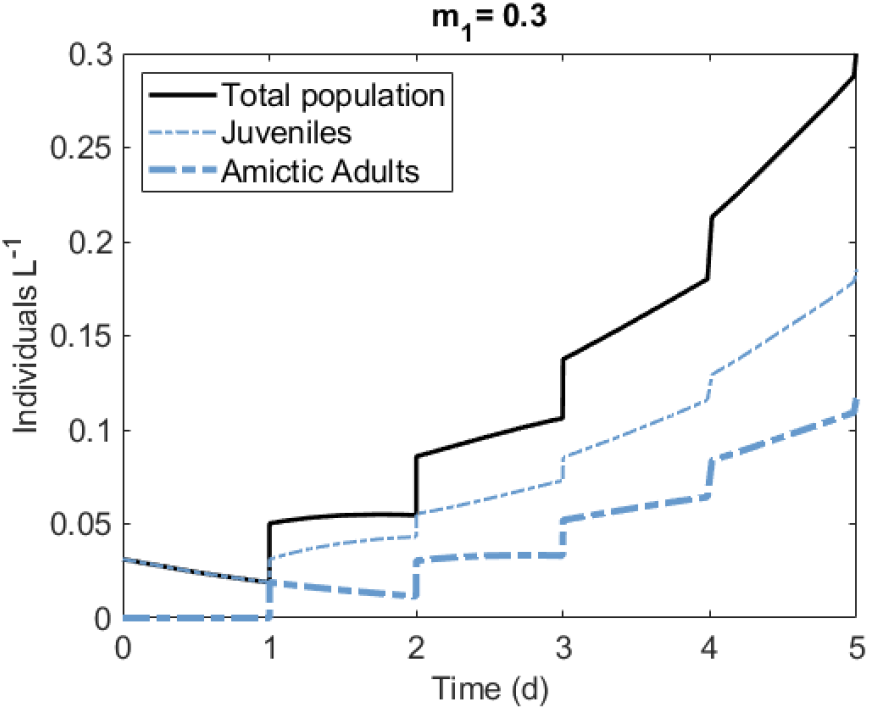
close-up of Fig. 3A highlighting the discontinuities in growth that result from hatching and maturing dynamics. At discrete intervals (in this case daily), new juveniles hatch and juveniles that have survived to adulthood mature.

**Figure S5:**
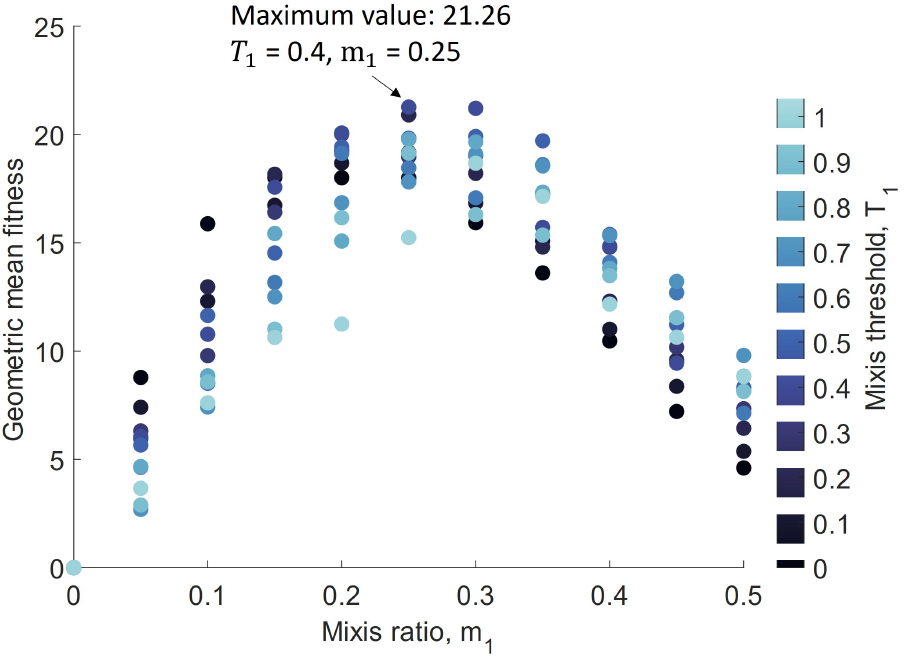
Fitness of monomorphic populations of phenotypes without a generational mictic block at a finer scale than Figure 4A. When *G*_1_ = 0, the strategy with the highest average fitness has a mixis ratio of 0.25 at a resolution of 0.05 and a mixis threshold density of 0.4 at a resolution 0.1.

**Figure S6:**
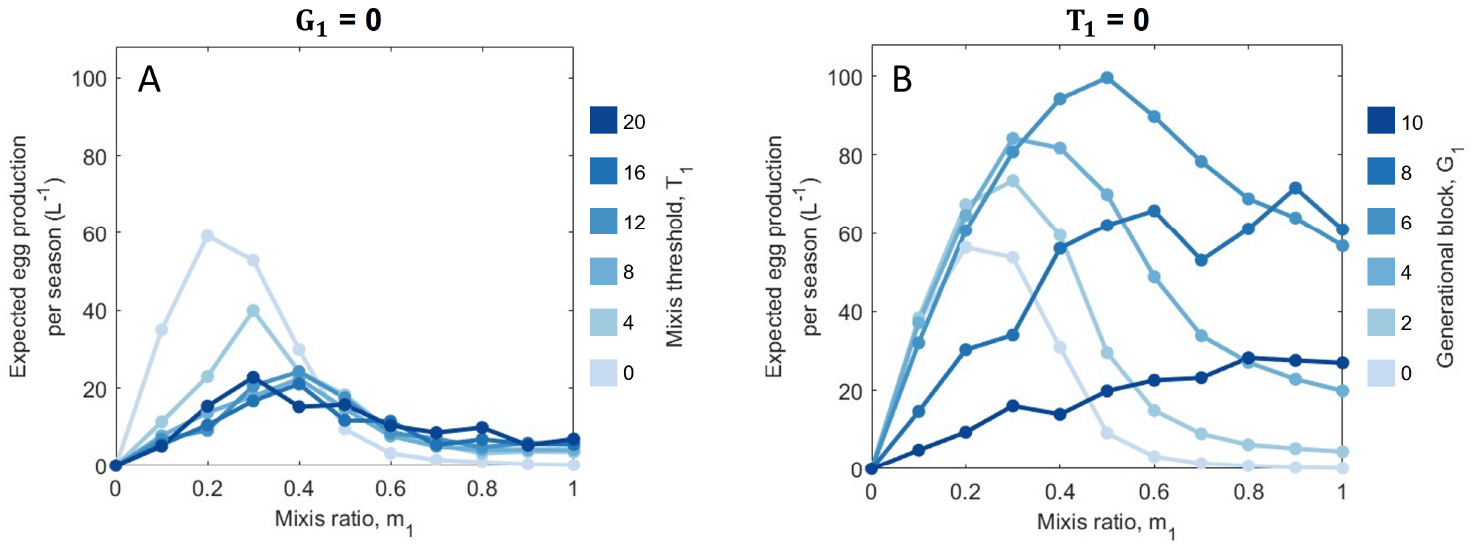
Expected egg production per season by monomorphic populations (A) without a generational mictic block and (B) without a mixis threshold density. Values plotted are the average eggs produced in 40 consecutive seasons of stochastic length (between 10 and 51 days) across 40 replicates with initial resting egg density equal to *B*_*max*_. All parameters not indicated within the figure have values shown in Table 1.

**Figure S7:**
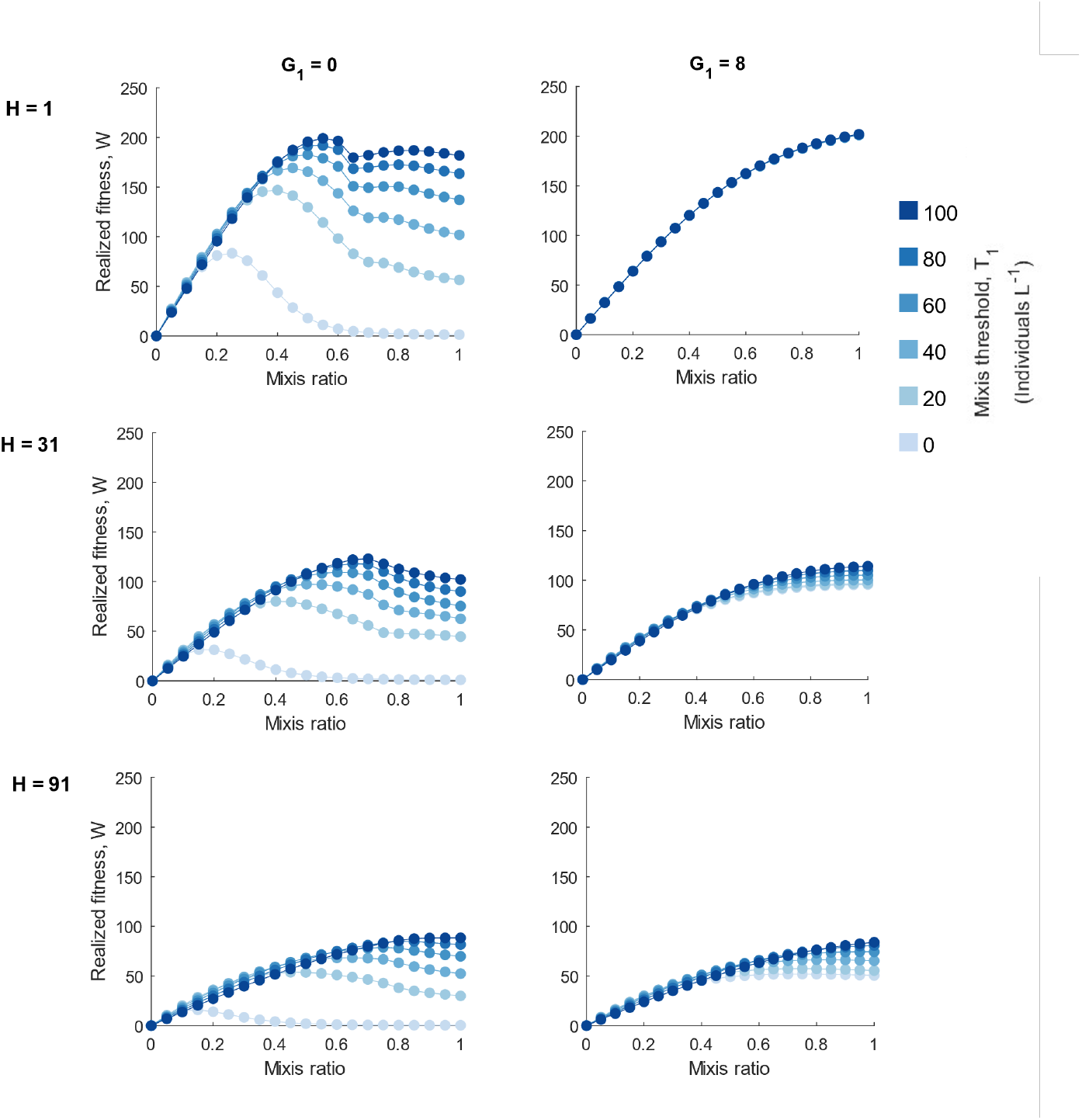
Realized fitness of monomorphic populations with multiple values of *H* (the end of the resting egg hatching period) indicated for each row. Seasons of length 30 days were simulated with all other parameter values indicated in Table 1 and initial resting egg density equal to *B*_*max*_ = 1. Populations that hatch earlier (low H), have similar fitness to standard populations in longer seasons (Fig. 5E), but trends across mixis phenotypes are unchanged.

**Figure S8:**
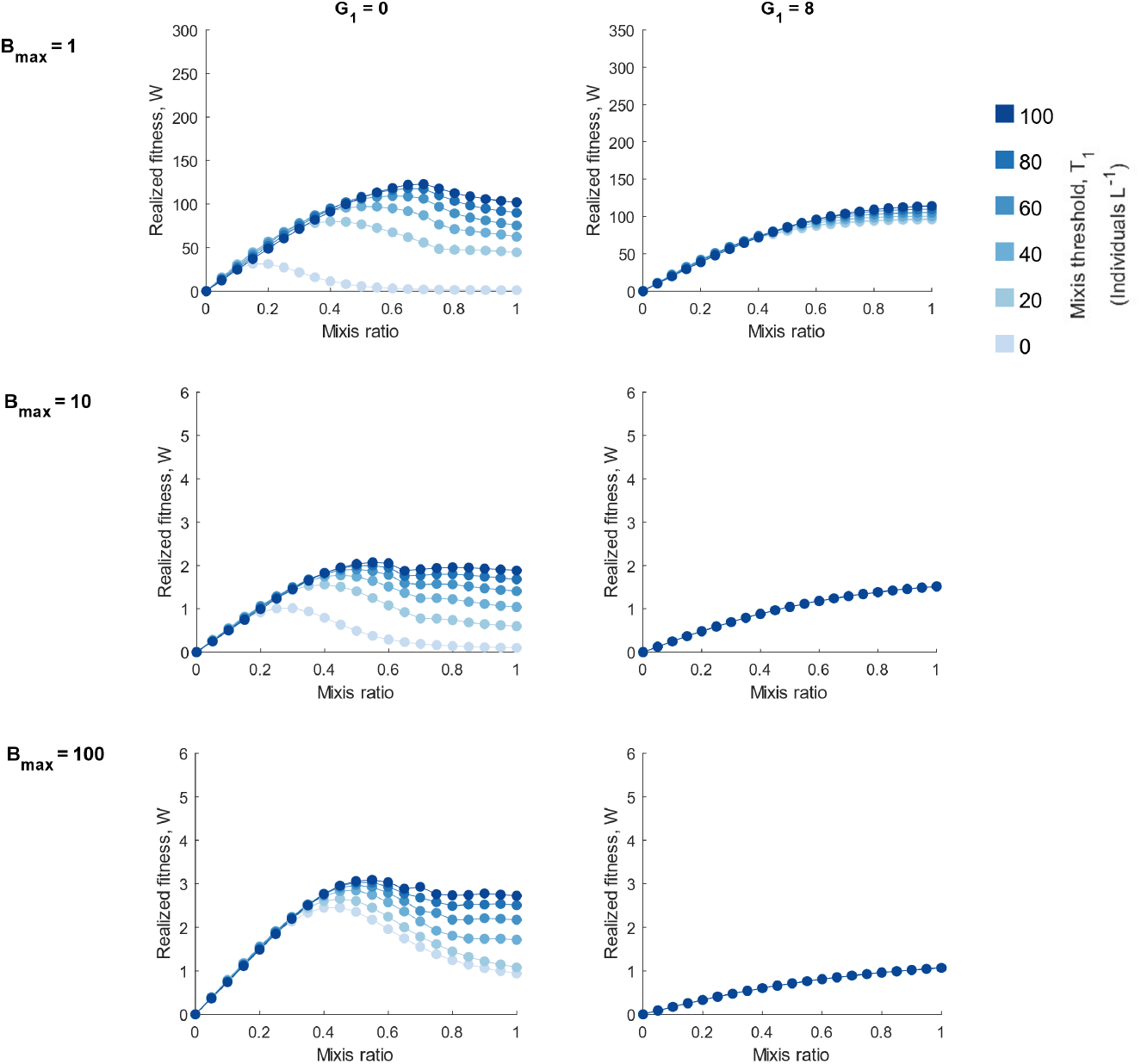
Realized fitness of monomorphic populations with multiple values of *B*_*max*_ (the initial size of the resting egg pool and maximum that can survive between seasons) indicated for each row. Seasons of length 30 days were simulated with all other parameter values indicated in Table 1.

**Figure S9:**
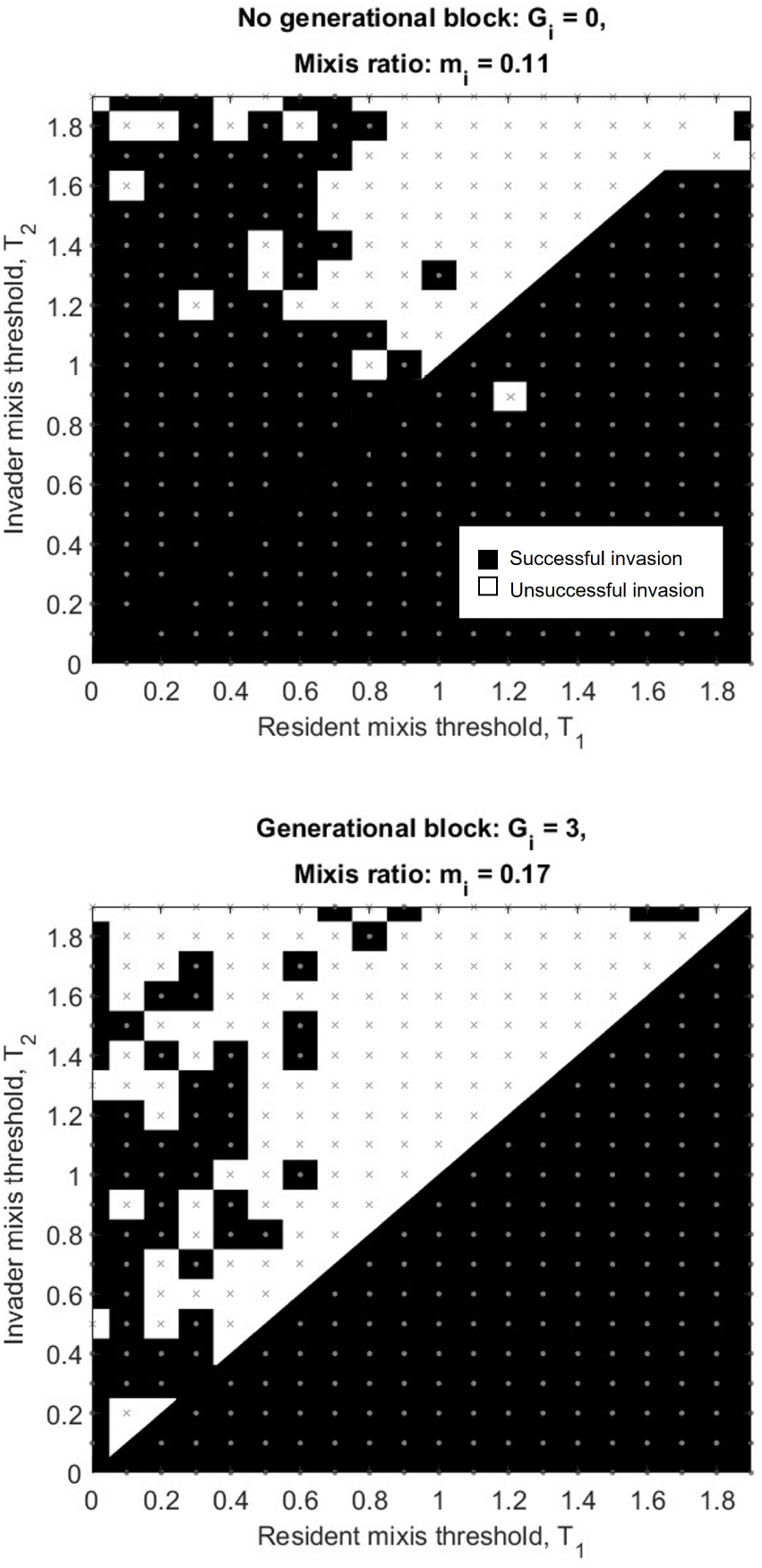
Pairwise invasibility plot for rotifer phenotypes with variable mixis threshold densities. As in Fig. 6, invasion was deemed successful if, out of 20 simulations of 40 growing seasons each, the invader frequency increased on average from its initial frequency of 0.05. White squares and cross marks indicate unsuccessful invasions, black sqaures and circle marks indicate successful invasion. Generally phenotypes with lower mixis threshold densities are more likely to successfully invade, and phenotypes with no mixis threshold density (*T*_*i*_ = 0) are able to invade all others. All parameters not indicated within the figure have values shown in Table 1

**Figure S10:**
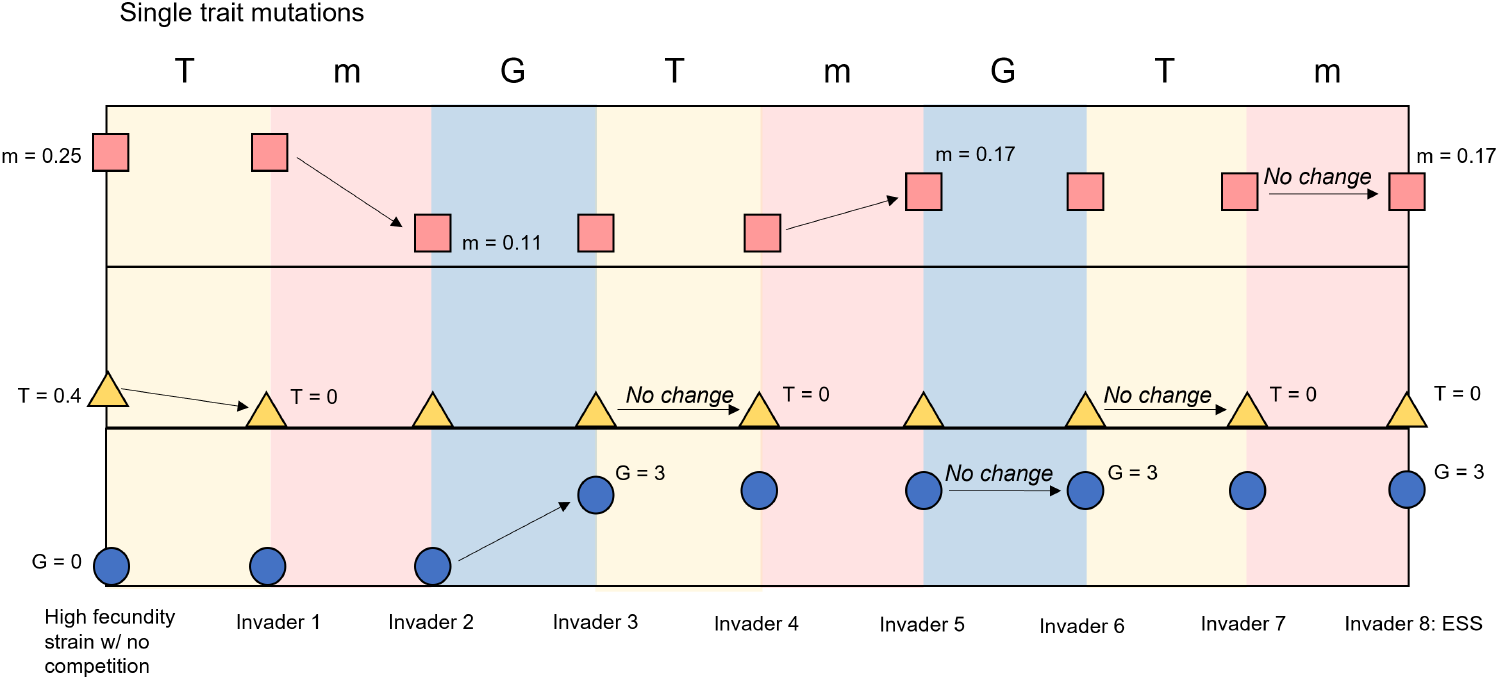
Schematic of sequential mutation simulation. We simulate a community evolving by allowing each dimension of the phenotype to change independently. For each single invasion step (columns), we construct a pairwise invasibility plot (Figs. S9,S11) and replace the resident with the invader that has a stable strategy along that single dimension. After five steps, the community reaches a phenotype that cannot be replaced by phenotypes with any single mutation.

**Figure S11:**
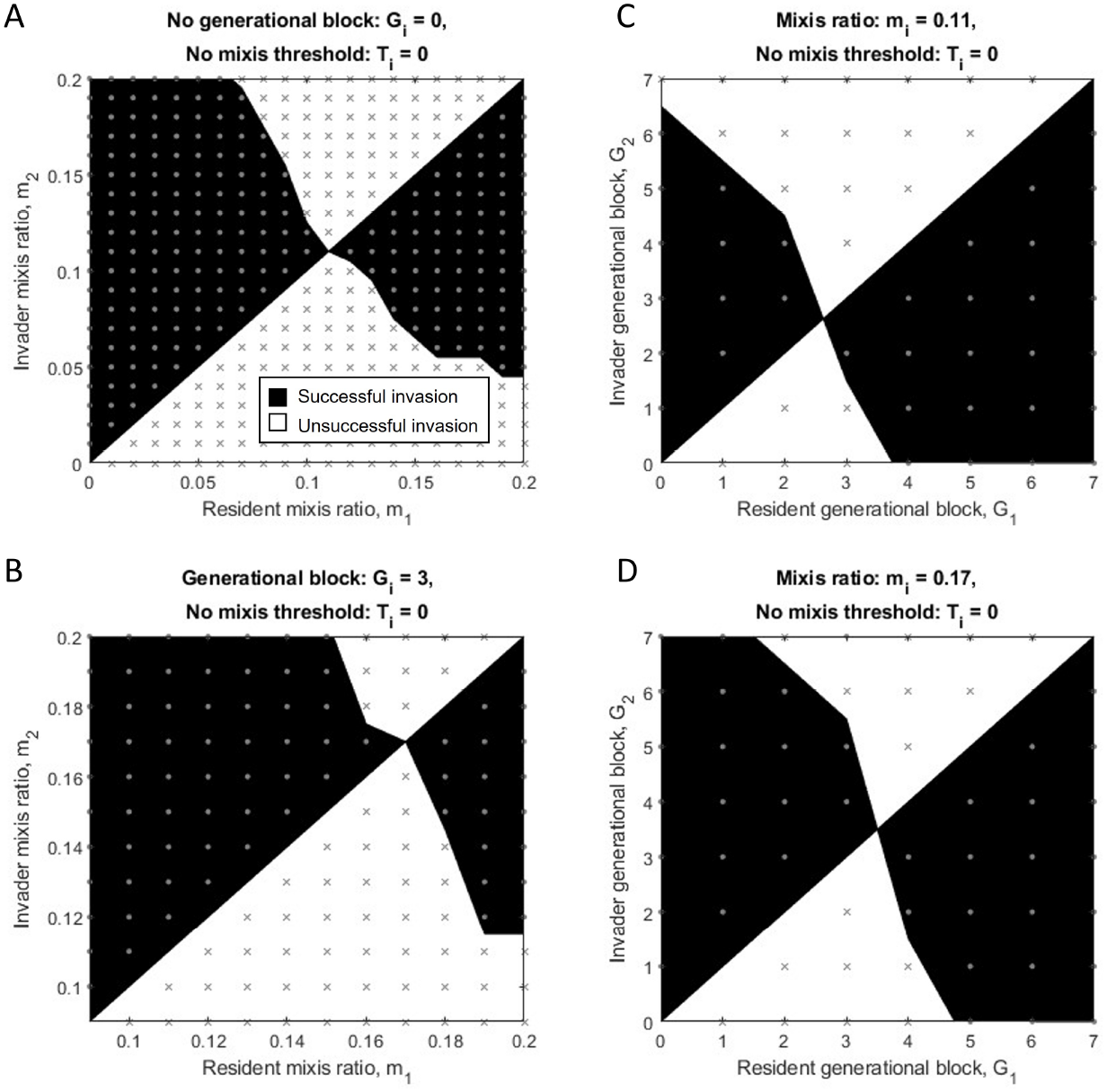
Pairwise invasibility plots for rotifers with A-B) variable mixis ratio and C-D) variable generational block. Panels A and C are exactly replicated from Fig. 6. As before, invasion was deemed successful if, out of 20 simulations of 40 growing seasons each, the invader frequency increased on average from its initial frequency of 0.05. Gray crosses indicate unsuccessful invasions; circles indicate successful invasions. Black and white regions are interpolated regions of success and failure, respectively. All parameters not indicated within the figure have values shown in Table 1.

**Figure S12:**
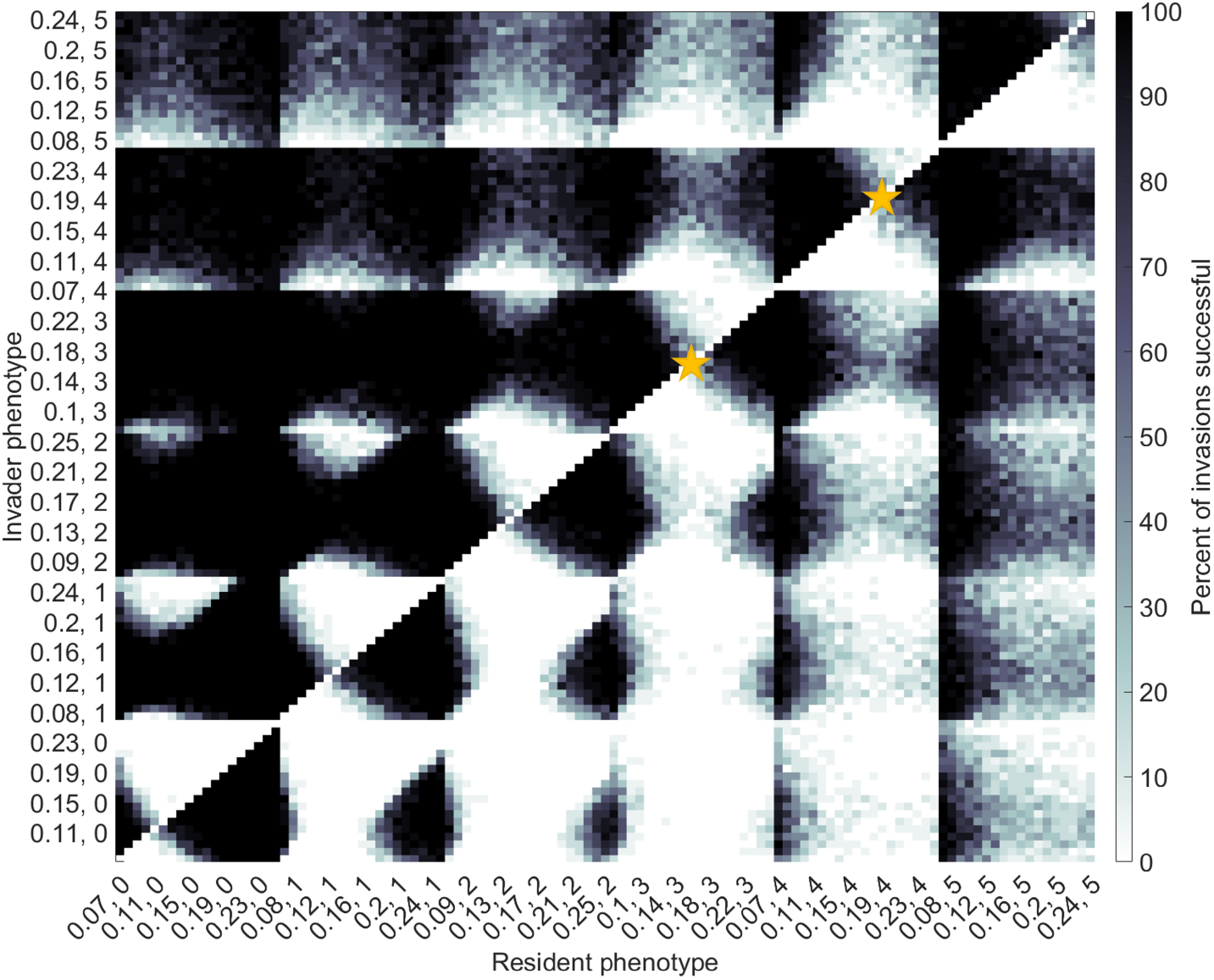
Pairwise invasibility plots for rotifers with variable mixis ratio and generational block length. Axes tick labels are values of *m*_*i*_ and *G*_*i*_ separated by a comma. Grid color shows percent of successful invasions out of 20 simulation experiments of 40 growing seasons each. Yellow stars indicate the two phenotypes that, on average, had highest invasion success and resistance to invasion by the other phenotypes within the entire grid shown.

**Figure S13:**
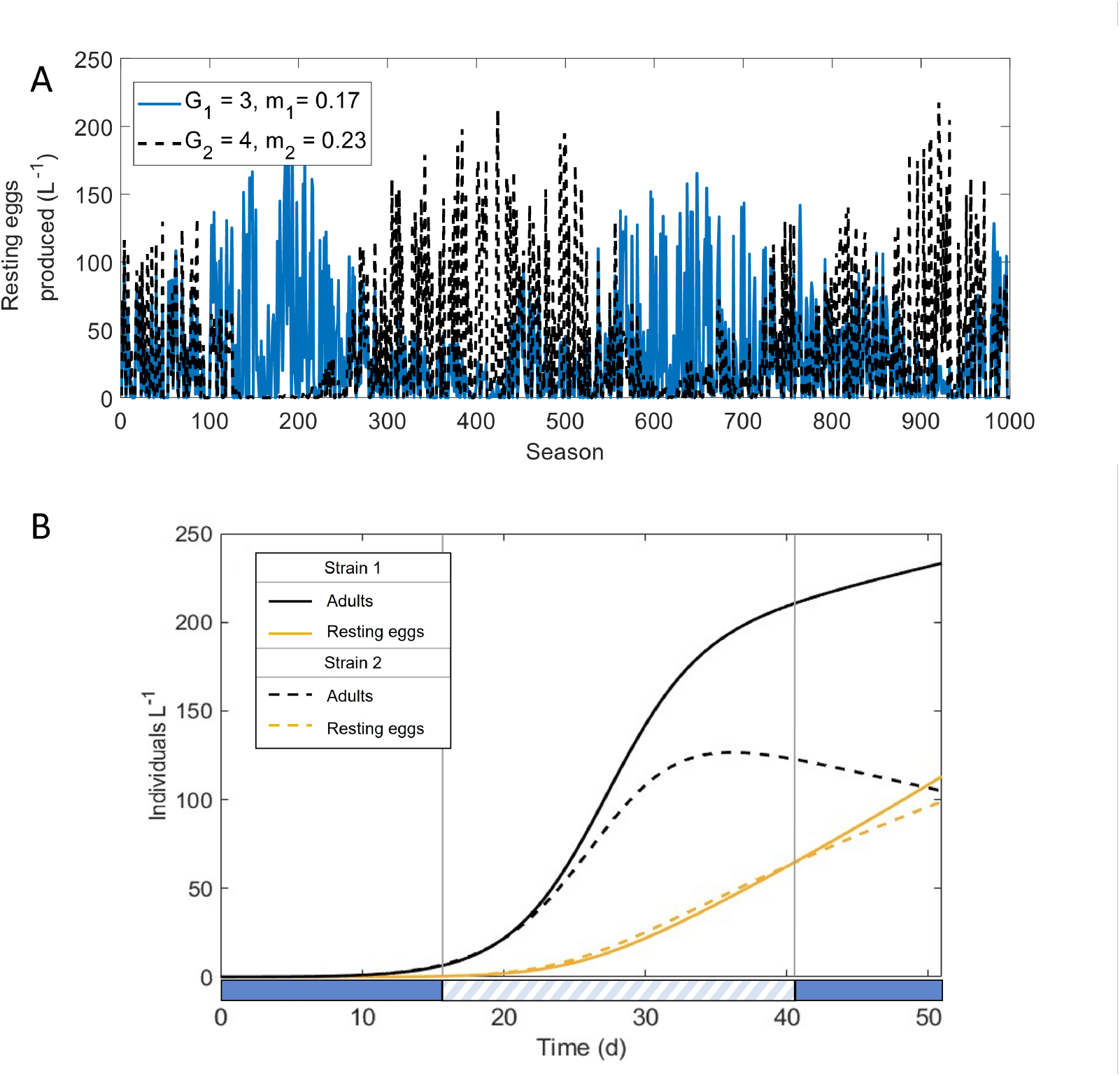
Population trajectories A) across seasons and B) within a single season for two phenotypes that can coexist within our standard environmental regime (uniformly distributed season lengths between 10 and 51 days). Solid lines represent the phenotype with shorter mictic block (*G*_1_ = 3) and lower mixis ratio (*m*_1_ = 0.17). Dashed lines represent a phenotype with longer mictic block (*G*_2_ = 4) and higher mixis ratio (*m*_2_ = 0.23). The abundance of the two phenotypes are equal 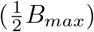 at the start of the simulation. Vertical gray lines and shaded bars along the x-axis indicate the two points in time when the density of resting eggs for the two strategies intersect. Mixis threshold density, *T*_*i*_, is 0 and all other parameter values in these simulations are those in Table 1.

**Figure S14:**
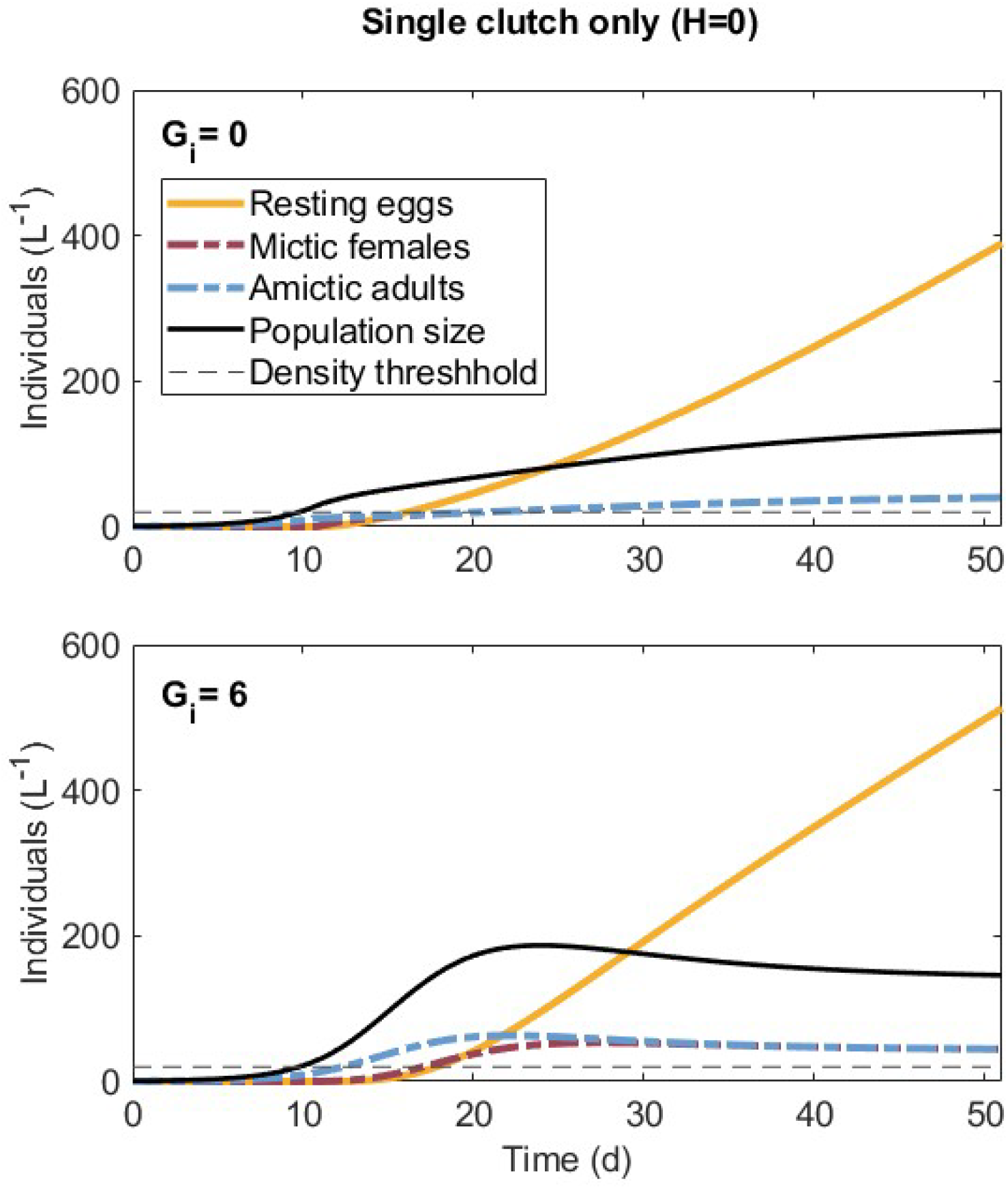
Population trajectories for monomorphic populations when all resting eggs hatch in a single clutch at t=0. Both phenotypes have a mixis ratio and threshold of 0.5 and 20 individual *L*^−1^, respectively. Phenotypes with *G*_*i*_ = 6 (bottom) produce more resting eggs in a long season than those with no generational block (top). Benefits of generational block are not limited to the growth of late hatching clones.

